# Differential GTP-dependent *in-vitro* polymerization of recombinant Physcomitrella FtsZ proteins

**DOI:** 10.1101/2024.02.14.580282

**Authors:** Stella W. L. Milferstaedt, Marie Joest, Lennard L. Bohlender, Sebastian N. W. Hoernstein, Buğra Özdemir, Eva L. Decker, Chris van der Does, Ralf Reski

**Author notes:** Euro-BioImaging Bio-Hub, EMBL, Meyerhofstraße 1, 69117 Heidelberg, Germany.

## Abstract

Bacterial cell division and plant chloroplast division require self-assembling Filamentous temperature-sensitive Z (FtsZ) proteins. FtsZ proteins are GTPases sharing structural and biochemical similarities with eukaryotic tubulin. In the moss Physcomitrella, the morphology of the FtsZ polymer networks varies between the different FtsZ isoforms. The underlying mechanism and foundation of the distinct networks is unknown. Here, we investigated the interaction of Physcomitrella FtsZ2-1 with FtsZ1 isoforms *via* co-immunoprecipitation and mass spectrometry, and found protein-protein interaction *in vivo*. We tagged FtsZ1-2 and FtsZ2-1 with different fluorophores and expressed both in *E. coli*, which led to the formation of defined structures within the cells and to an influence on bacterial cell division and morphology. Furthermore, we have optimized the purification protocols for FtsZ1-2 and FtsZ2-1 expressed in *E. coli* and characterized their GTPase activity and polymerization *in vitro*. Both FtsZ isoforms showed GTPase activity. Stoichiometric mixing of both proteins led to a significantly increased GTPase activity, indicating a synergistic interaction between them. In light scattering assays, we observed GTP-dependent assembly of FtsZ1-2 and of FtsZ2-1 in a protein concentration dependent manner. Stoichiometric mixing of both proteins resulted in significantly faster polymerization, again indicating a synergistic interaction between them. Under the same conditions used for GTPase and light scattering assays both FtsZ isoforms formed filaments in a GTP-dependent manner as visualized by transmission electron microscopy (TEM). Taken together, our results reveal that Physcomitrella FtsZ1-2 and FtsZ2-1 are functionally different, can synergistically interact *in vivo* and *in vitro*, and differ in their properties from FtsZ proteins from bacteria, archaea and vascular plants.

## 2. Introduction

Plants harvest energy from oxygenic photosynthesis, during which light is the energy source for the fixation of atmospheric CO_2_ in organic compounds, while O_2_ is produced as a side product. In eukaryotic cells, photosynthesis depends on chloroplasts [1]. These cell organelles derived from endosymbiosis, where a eukaryote engulfed a free-living, photoautotrophic cyanobacterial-like prokaryote [2, 3, 4]. While some genes of prokaryotic origin disappeared over time, or were retained in the organellar genome, respectively, most were transferred to the host nucleus after the endosymbiotic event [5]. The relation of these organelles to prokaryotes explains why chloroplasts do not develop *de novo* but divide by binary fission [6], while at the same time suggesting that prokaryotic cytokinesis and cell organelle division are conserved mechanisms [7, 8].

Most bacteria and chloroplasts of bryophytes, comprising mosses, liverworts, and hornworts, are surrounded by a peptidoglycan wall [9, 10], which needs to be remodelled during bacterial cell division [11, 12]. Proteins involved in bacterial cell division are also responsible for plastid division in plants [6, 8]. Both mechanisms require Filamentous temperature-sensitive Z (FtsZ) proteins as a key component of the division machinery [13, 14]. FtsZ proteins were named after one of the filamentous temperature-sensitive *Escherichia coli* (*E. coli*) mutants, which grows filamentous due to their inability to divide at elevated temperatures [15]. FtsZ proteins assemble into a ring structure, the Z-ring, thus determining the future division site of bacteria and chloroplasts [16, 17, 18]. In *E. coli* and *Bacillus subtilis*, FtsZ-function is coupled to peptidoglycan synthesis [11, 19]. Remarkably, bacterial cell division as well as division of moss chloroplasts, but not plastid division of vascular plants, is sensitive to beta-lactam antibiotics [20].

FtsZ proteins are soluble guanosine triphosphatases (GTPases) [21, 22, 23] that share structural and biochemical similarities with the eukaryotic cytoskeletal protein tubulin [24, 25]. FtsZ proteins have similar functions in bacteria and plastids [7, 8]. The nuclear-encoded plant FtsZ proteins are imported into the chloroplast [7], where they are required for plastid division as well as plastid shaping [26]. In most bacteria, FtsZ is encoded by a single gene [27, 28], while plants and archaea, except the TACK and ASGARD superphyla, encode predominantly two FtsZ isoforms [25, 29]. For instance, *Arabidopsis thaliana* (Arabidopsis) encodes FtsZ1 and FtsZ2 [7, 14]. These arose from a gene duplication [29], in which plant FtsZ1 arose from FtsZ2 [31, 32].

In the vascular plant Arabidopsis, both FtsZ isoforms have non-redundant functions since their independent expression inhibition led in both cases to a reduced number of chloroplasts [14]. The rings formed by Arabidopsis FtsZ1 and FtsZ2 colocalize *in vivo* at the chloroplast centre [17] and form heteropolymers at the future division site upon recombinant expression in the yeast *Pichia pastoris* (new species name *Komagataella phaffii*) [18]. Also, the polymerization behaviour of Arabidopsis FtsZ isoforms has been characterized *in vitro* [33, 34]. Here, FtsZ1 and FtsZ2 show differences in their GTPase activity and polymerization mechanism and only FtsZ2 polymerized on its own [34].

Furthermore, FtsZ1 and FtsZ2 show differences regarding their protein structure. In general, FtsZ proteins contain two domains, a widely conserved N-terminal GTPase domain and a C-terminal region [32, 35]. The GTPase domain comprises a GTP-binding domain and a GTPase-activating domain. Together, they represent the core structure of FtsZ [18], since the C-terminal region is more variable [36]. The N-terminal domain is responsible for GTP binding and hydrolysis [35]. FtsZ2 proteins contain an additional short motif at the C-terminus that is missing in FtsZ1 [31, 32]. In Arabidopsis, this motif is responsible for the interaction of FtsZ2 with transmembrane proteins [32]. Recently, a conserved sequence motif of the C-terminal domain of Arabidopsis FtsZ1 was identified that is involved in membrane binding and interactions with other proteins of the plastid division machinery [37]. Moreover, the N-as well as the C-terminus of both isoforms are involved in polymer formation of FtsZ1 and FtsZ2 [38].

The moss Physcomitrella (new species name *Physcomitrium patens*) encodes the high number of five different FtsZ proteins (FtsZ1-1, FtsZ1-2, FtsZ2-1, FtsZ2-2, FtsZ3), which fall in three clades (FtsZ1, FtsZ2, FtsZ3), indicating neofunctionalisation of the different isoforms during evolution [30, 39, 40]. Arabidopsis FtsZ1 and FtsZ2 are orthologues to Physcomitrella FtsZ1 and FtsZ2 [39, 40]. Among those FtsZ isoforms, FtsZ2-1 was proven to be involved in chloroplast division [8, 41, 42].

The Physcomitrella FtsZ proteins assemble in network-like structures inside the chloroplast [26, 43]. Using reverse genetics, the diverse and non-redundant functions of the five FtsZ isoforms in Physcomitrella were investigated [42]. Here, the analysis of distinct single and double knockout mutants of Physcomitrella *ftsZ* genes revealed their roles in chloroplast division, chloroplast shaping, as well as their network assembling properties within the chloroplast [26, 42]. Due to these characteristics, the Physcomitrella FtsZ proteins ensure plastid stability and structural integrity [26, 44]. Since this FtsZ network in moss plastids resembles the structure of the eukaryotic cytoskeleton, the term ‘plastoskeleton’ was coined to describe its organization and function in plastids [26, 45]. Subsequently, it was discovered that not only eukaryotes but also bacteria have a highly dynamic cytoskeleton at which FtsZ is a part [46–49].

Unlike described for other plants, proteins of the nuclear *ftsZ* genes are not only imported into plastids but are dually targeted to chloroplasts and the cytosol in Physcomitrella, suggesting that they have functions beyond plastid division [50]. Interestingly, this coincides with a low diversification of the cytoskeletal protein family tubulin in Physcomitrella [51]. Subsequently, Physcomitrella FtsZ proteins were found to be involved in cell patterning, plant development, and gravity sensing [42]. Additionally, distinct localization-dependent *in-vivo* interactions among four of the five Physcomitrella FtsZ isoforms were revealed by using fluorescence resonance energy transfer (FRET) [52].

The morphology of the Physcomitrella FtsZ polymer networks varies between the different isoforms. *In-vivo* network analysis using GFP-tagged FtsZ1-2 and FtsZ2-1 indicated functional differences between them [53]. While the FtsZ2-1 network is exclusively formed within the chloroplast, FtsZ1-2 networks form long, extra-plastidic extensions and may play a role in the formation of stromules [53], stroma-filled, tubular connections between chloroplasts [54]. The FtsZ1-2 network also contains significantly more nodes than the FtsZ2-1 network, while so-called meganodes (extraordinarily large nodes) occur only with FtsZ2-1. Moreover, FtsZ2-1 filaments are more curved *in vivo*, thicker in segment size and resemble the microtubules of the cytoskeleton more than FtsZ1-2. This resemblance hints towards mechanical properties that might be interesting when using Physcomitrella FtsZ filaments to develop sustainable material systems [55]. Based on the identified morphological differences of Physcomitrella FtsZ1-2 and FtsZ2-1 [53], the mechanical behaviour and load-bearing responses of the two Physcomitrella FtsZ isoforms were investigated in *in-silico* experiments, relating the structure to the function of the FtsZ protein networks [55–57]. The capability of the plastoskeleton to maintain its load-bearing structure even under distortion is based on its material properties as polymers and more importantly, on the structural features of the network [55].

Besides the mechanical properties of the FtsZ isoforms, knowledge about the polymer-forming properties of Physcomitrella FtsZ proteins as a GTP-driven structure is of particular interest. Combining the data about the load-bearing structure of the plastoskeleton with knowledge about the biochemical foundations could pave the way for adaptive biomimetic materials [58] or 3D-printed biopolymer materials for tissue engineering [59]. In various tissues, Physcomitrella FtsZ1-2 and FtsZ2-1 have relatively high expression levels, suggesting division-independent, stable networks within the chloroplast [53]. We examined GTPase activity, polymer assembly, and formation of the Physcomitrella FtsZ network *in vitro*. We expressed Physcomitrella FtsZ1-2 and FtsZ2-1 in *E. coli*, purified the proteins, and analysed their GTP-dependent polymerization individually and together. Furthermore, we investigated the interaction of these isoforms *in vivo*, and visualised both FtsZ isoforms in *E. coli* cells individually and in combination without potential interaction partners.

Different bacterial FtsZ (e.g., *B. subtilis* FtsZ and *E. coli* FtsZ [60], *Agrobacterium tumefaciens* FtsZ [61], *Synechococcus elongatus* [23], archaeal FtsZ (*Haloferax volcanii* [62, 63]), and plant FtsZ (Arabidopsis [23, 34]) were already investigated regarding their assembly and biochemical properties. Here, we provide insights into the GTP-dependent polymerization and the assembly properties of FtsZ1-2 and FtsZ2-1 of the moss Physcomitrella.

## 3. Results

### 3.1. *In-vivo* interaction of Physcomitrella FtsZ2-1 with FtsZ1

Distinct localization-dependent interactions among four of the five Physcomitrella FtsZ isoforms have been revealed by overexpression as GFP fusions [52]. Here, we independently reassessed these results on a physiological level with special focus on the interactions between FtsZ1 isoforms and FtsZ2-1.

To address this, an in-frame fusion of Physcomitrella FtsZ2-1 with a single linker-GFP (Figure 1A, Supplemental Figure S1A) at the endogenous locus was generated *via* targeted knock-in (Supplemental Table S1), employing the highly efficient homologous recombination in this moss [64, 65]. Twenty-one selected lines were screened by PCR for the presence of the GFP coding sequence (CDS). From these, sixteen positive candidate lines were selected and the correct integration of the knock-in construct was analysed by PCR (Supplemental Figure S1A, B). From this analysis three of the fifteen positive lines were chosen for a test co-immunoprecipitation (Co-IP, Supplemental Figure S1C). Finally, line #513 was chosen for quantitative Co-immunoprecipitations (Co-IP), performed using GFP-Trap Magnetic Particles, with Physcomitrella wild type (WT) as a negative control. Label-free quantitation values (LFQ) obtained from a *MaxQuant* search [66, 67] were used and significant interacting partners were identified at a false discovery rate (FDR) of 1 % (Figure 1B). The significant interacting proteins can be found in Supplemental Table S2.

**Figure 1.**
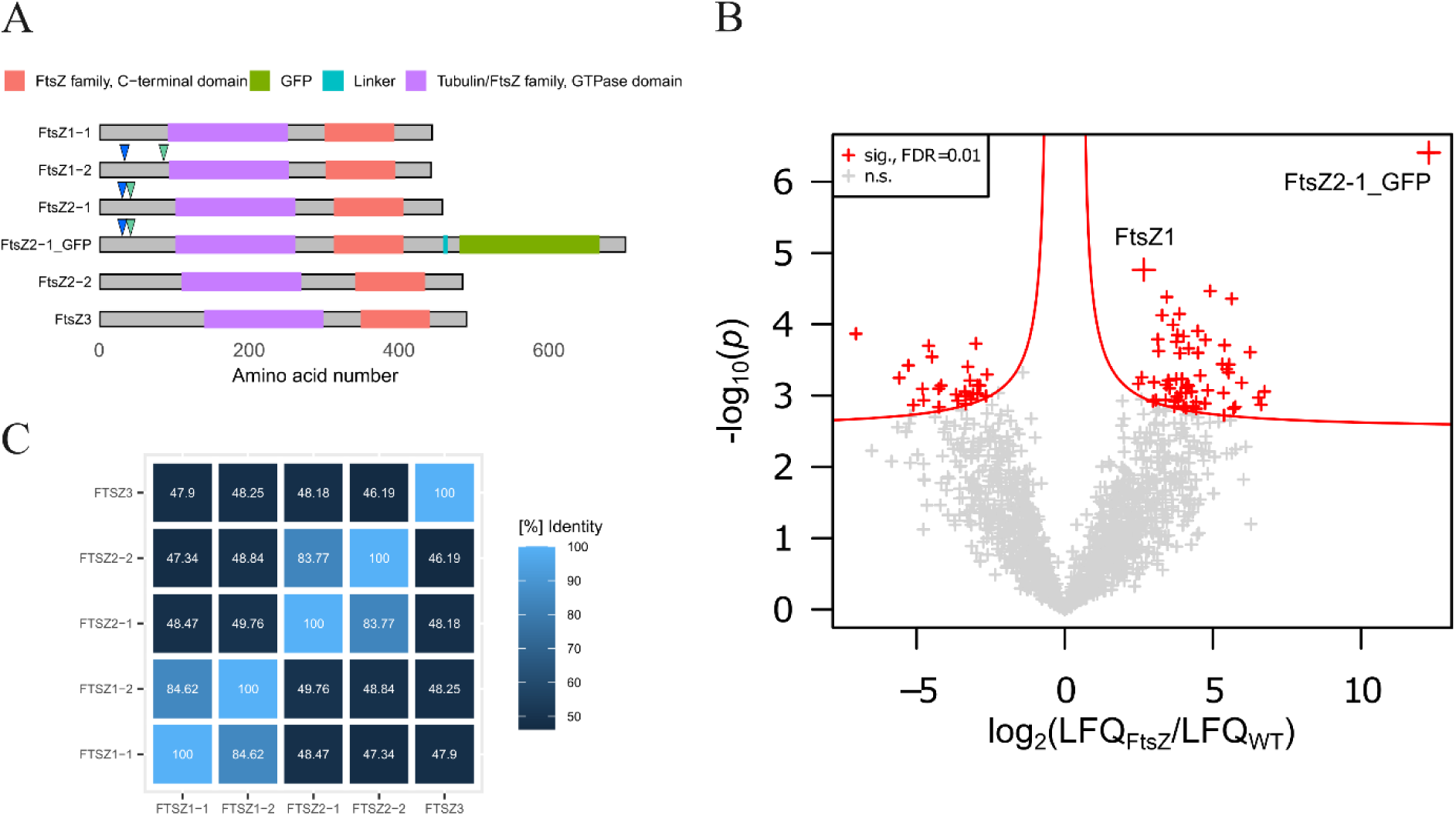
FtsZ domain structures, sequence identity and analysis of the co-immunoprecipitations (Co-IPs) against GFP-tagged FtsZ2-1 protein. **(A)** Domain structures of the five Physcomitrella FtsZ proteins and the FtsZ2-1_GFP fusion protein. Depicted are PFAM [68] domains (FtsZ family, C-terminal domain: PF12327; Tubulin/FtsZ family, GTPase domain: PF00091; GFP: PF01353) and the poly-G linker in the FtsZ2-1_GFP fusion protein. Arrows indicate predicted chloroplast transit peptide (cTP) cleavage sites (blue: ChloroP1.1; green: TargetP2.0). Predicted cTP cleavage sites of other FtsZ isoforms than FtsZ1-2 and FtsZ2-1 are not shown. The image was created with the *R* package *drawProteins* [69]. (**B**) The Volcano plot shows the log_2_ ratios of normalized LFQ (label-free quantitation) intensities plotted against log_10_ of adjusted p-values. Proteins significantly enriched in the GFP-tagged pulldown are shown as red crosses with a false discovery rate (FDR) of 0.01 %. Physcomitrella wild type (WT) served as a negative control. Proteins not significant for neither WT nor FtsZ2-1-GFP are visualized as grey crosses. The Co-IP was performed with GFP-Trap Magnetic Particles M-270 (ChromoTek GmbH, Planegg, Germany) and Physcomitrella protonema tissue homogenized from suspension culture. The isoforms of FtsZ1 (FtsZ1-1, FtsZ1-2) could not be distinguished on the basis of the peptides identified in this approach and are thus grouped (FtsZ1). (**C**) Matrix showing the sequence identity ([%]) between the five different FtsZ isoforms. Identity values were obtained from a multiple sequence alignment using protein sequences done with the UniProt *Align* tool (https://www.uniprot.org/align).

From this, Physcomitrella FtsZ1 isoforms were identified as significant interactors of FtsZ2-1 (Figure 1). Thus, we confirmed that FtsZ2-1 is interacting with FtsZ1 proteins *in vivo*. However, due to their high sequence similarity (Figure 1C) and the limited number of matching peptides, the isoforms FtsZ1-1 and FtsZ1-2 could not be distinguished in these experiments. Hence, we were able to confirm that Physcomitrella FtsZ2-1 is interacting with Physcomitrella FtsZ1 proteins *in vivo*, but we could not explicitly narrow this down to an interaction with FtsZ1-2. As CO-IPs do not allow to assess spatial information of interactions, they can be direct or indirect.

### 3.2. Expression of Physcomitrella FtsZ1-2 and FtsZ2-1 in *E. coli*

The respective plastoskeletal morphologies of Physcomitrella FtsZ1-2 and FtsZ2-1, which were elucidated before [53], hint towards functional differences of these two isoforms. Therefore, GTPase activity, polymer assembly, and the formation of Physcomitrella FtsZ1-2 and FtsZ2-1 filaments *in vitro* were studied.

In order to produce Physcomitrella FtsZ1-2 and FtsZ2-1 in *E. coli*, the coding sequences (CDS) of both FtsZ isoforms were optimized for codon usage in bacteria (Supplemental Figures S2 and S3). In total, 304 bases were changed for FtsZ1-2 and 357 bases were changed for FtsZ2-1. The GC content was concurrently increased from 49 % to 51 % for FtsZ1-2, and from 49 % to 52 % for FtsZ2-1. The *in-silico* predicted N-terminal transit peptides for Physcomitrella FtsZ1-2 (Pp3c19_2490) and Physcomitrella FtsZ2-1 (Pp3c11_17860) were 34 aa and 31 aa (ChloroP 1.1, http://www.cbs.dtu.dk/services/ChloroP/), respectively. The predictions of TargetP-2.0 (http://www.cbs.dtu.dk/services/TargetP/) and ChloroP 1.1 differ (Table 1, Figure 1A) and the exact estimation of cTPs remains challenging. The shorter predictions by ChloroP 1.1 were chosen so as not to affect the functional domains of both proteins (Figure 1A).

**Table 1.**
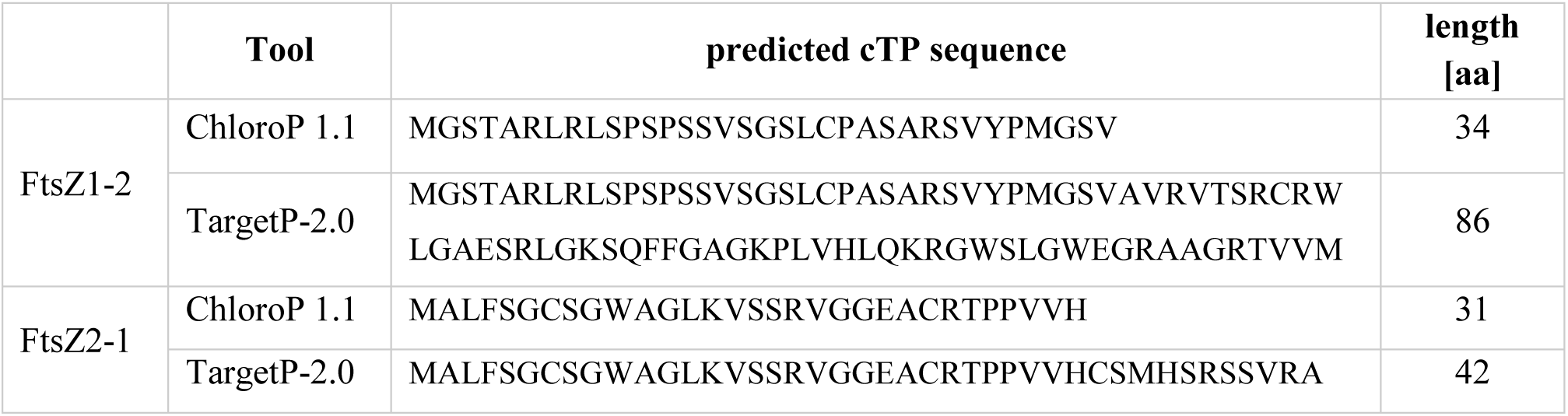
Comparison of sequence and length of the chloroplast transit peptide (cTP) for Physcomitrella FtsZ1-2 and FtsZ2-1 predicted by ChloroP 1.1 and TargetP-2.0. Length of cTP in amino acids (aa).

The respective codon-optimized FtsZ1-2 and FtsZ2-1 sequence, fused to a C-terminal 8 x His-tag encoding sequence, were subcloned into the pLATE11 expression vector (Supplemental Tables S3 and S4) and with the correct plasmids subsequently transformed into BL21 Star™ (DE3) cells. Immunoblots of IPTG induced strains confirmed the expression of both isoforms (Supplemental Figure S4).

### 3.3. Physcomitrella FtsZ proteins localize in foci in elongated *E. coli* cells

To investigate the behaviour of each isoform individually and in combination within a heterologous system, fluorescent FtsZ fusions were employed. *E. coli* FtsZ, which is non-functional in Physcomitrella and does not interact with Physcomitrella FtsZ isoforms in FRET experiments [51], is not anticipated to interact with Physcomitrella proteins. FtsZ1-2 was fused at the N-terminus with enhanced Green Fluorescent Protein (eGFP), while FtsZ2-1 was fused at the N-terminus with mKusabira-Orange2 (mKOrange2, mKO2). To serve as controls, eGFP-His and mKO2-His were used respectively. All CDS were cloned into the inducible pLATE11 expression vector. For the co-expression of eGFP-FtsZ1-2 and mKO2-FtsZ2-1, the latter expression cassette was introduced into the initially generated pLATE11_eGFP-FtsZ1-2 vector. BL21 Star™ (DE3) cells were transformed with the final plasmids. Confocal microscopy of IPTG-induced bacteria revealed that the eGFP-His control was localized in a diffuse pattern within the cell (Figure 2A). The cells expressing eGFP-His grew normally and were approximately 1-2 µm long. The eGFP-FtsZ1-2 protein was localized in numerous foci within the cell (Figure 2B), and, in some instances, formed single filaments or filamentous networks (Figure 3, Supplemental Figures S5A, B, E, and S7C). Cells expressing eGFP-FtsZ1-2 were longer than those expressing eGFP-His, showing a length distribution between 5 and 10 µm (Figure 2B, and Supplemental Figure S5). The distribution of the eGFP-FtsZ1-2 signal suggests the assembly of functional structures, which is further supported by the filamentous phenotype of the *E coli* cells expressing eGFP-FtsZ1-2, which resembles the phenotype of *E. coli* strains overexpressing FtsZ or *fts*Z depletion strains.

**Figure 2.**
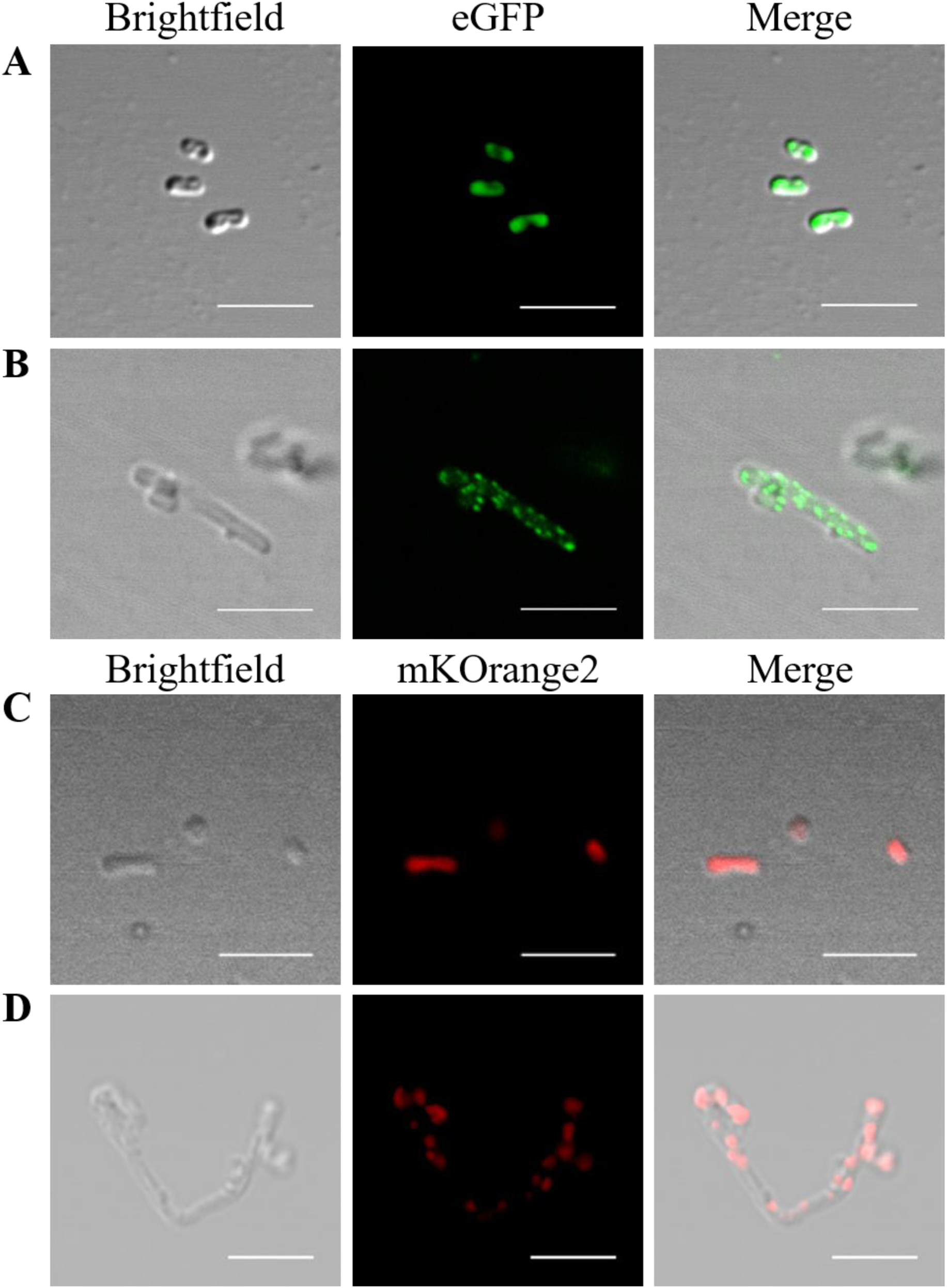
Cellular positioning of Physcomitrella FtsZ1-2 and FtsZ2-1 fluorescent fusion proteins in transformed *E. coli* cells. Confocal fluorescence microscopy of (**A**) eGFP-His, (**B**) eGFP-tagged Physcomitrella FtsZ1-2, (**C**) mKOrange2-His and (D) mKOrange2-tagged Physcomitrella FtsZ2-1 in *E. coli.* Scale bars 5 µm.

**Figure 3.**
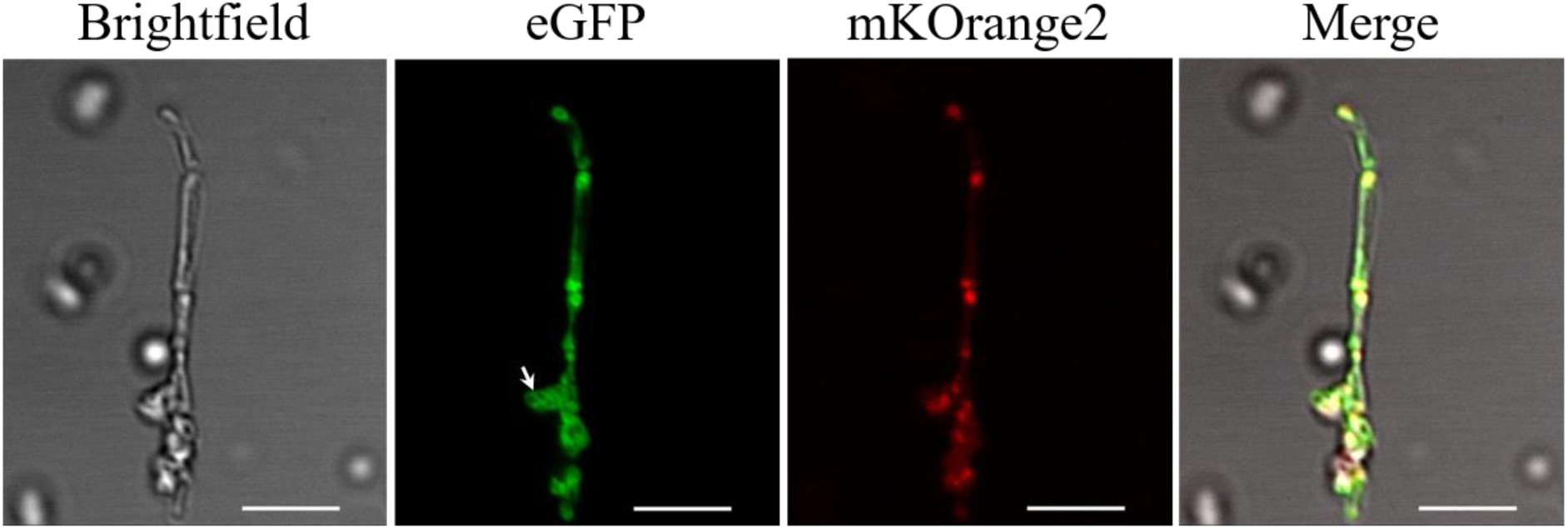
Cellular positioning of combined expression of Physcomitrella FtsZ1-2 and FtsZ2-1 fluorescent fusion proteins in *E. coli*. Confocal fluorescence microscopy of *E. coli* cells co-expressing eGFP-tagged Physcomitrella FtsZ1-2 and mKOrange2-tagged Physcomitrella FtsZ2-1. Filamentous eGFP-FtsZ1-2-derived network formation is highlighted with an arrow. Scale bars 5 µm.

As with eGFP-His, the mKO2-His control exhibited a diffuse localization throughout the cell (Figure 2C). Cells transformed with mKO2-His also grew normally and were approximately 1-2 µm long. The mKO2-FtsZ2-1 protein was distributed in foci throughout the whole cell. The cells were elongated, ranging between 5 and 20 µm in length (Figure 2D, Supplemental Figure S6). In analogy to eGFP-FtsZ1-2, we infer that mKO2-FtsZ2-1 also assembles in functional structures when expressed in *E. coli* cells.

In general, the signal of mKO2-FtsZ2-1 expressed in *E. coli* was weaker than the signal of eGFP-FtsZ1-2. Furthermore, it was observed that the foci of the mKO2-FtsZ2-1 were less frequent than in eGFP-FtsZ1-2 and were situated at regular intervals inside the cell. In contrast to eGFP-FtsZ1-2, no filaments between the foci of mKO2-FtsZ2-1 were observed.

Confocal microscopy of *E. coli* cells containing the double construct eGFP-FtsZ1-2_mKO2-FtsZ2-1 confirmed the heterologous expression of eGFP-FtsZ1-2 and mKO2-FtsZ2-1 in *E. coli* (Figure 3). Cells expressing eGFP-FtsZ1-2_mKO2-FtsZ2-1 were elongated and ranged between 5 and 20 µm in length (Figure 3, Supplemental Figure S7). In these cells, eGFP-FtsZ1-2 was once again localized in foci and filamentous networks. Correspondingly, the distribution pattern of mKO2-FtsZ2-1 was similar as observed in the cells transformed with the single constructs: mKO2-FtsZ2-1 was distributed in foci throughout the whole cell. Remarkably, it was observed that eGFP-FtsZ1-2 and mKO2-FtsZ2-1 co-localize in these foci within the cell (Figure 3) suggesting a direct or indirect interaction between the two proteins. Furthermore, it was observed that bacterial cells expressing eGFP-FtsZ1-2, mKO2-FtsZ2-1 and the construct with both FtsZ fusion proteins together exhibited highly aberrant phenotypes (Figure 2, Figure 3, Supplemental Figures S5, S6 and S7), suggesting that expression of the two moss FtsZ proteins affected bacterial cell division and shaping.

### 3.4. Optimized purification of recombinant Physcomitrella FtsZ proteins

To investigate whether Physcomitrella FtsZ1-2 and FtsZ2-1 can form plastoskeleton-like structures in the absence of other proteins, we aimed to purify both proteins recombinantly produced in *E. coli*. Overexpression and initial attempts to purify the C-terminally His-tagged Physcomitrella FtsZ1-2 and FtsZ2-1 from *E. coli* revealed that both proteins were partially localized in inclusion bodies. Purification of the soluble fractions using standard affinity purification protocols resulted in relatively low yields and highly contaminated protein samples. Consequently, we sought to optimize the expression strategies for both FtsZ isoforms. To enhance the stability of the FtsZ proteins and reduce inclusion body formation, Physcomitrella FtsZ1-2 and FtsZ2-1, lacking the N-terminal transit peptides of 86 and 42 amino acids, respectively (Table 1), were fused to an N-terminal His6-SUMO tag, which was cleaved off after the first purification step. Various expression conditions were tested, with the highest amounts of soluble protein obtained after growth in a simplified version of autoinduction medium at 25 °C, as described by [70]. The buffer composition used during the purification process was subsequently optimized. Both proteins exhibited significant aggregation at lower and neutral pH values, as well as at higher salt concentrations. Stabilization was achieved in the presence of GDP and glycerol, whereas the presence of sodium inhibited activity after purification. The optimized purification buffer compositions were as follows: for FtsZ1-2, 50 mM glycine (pH 9.5), 150 mM KCl, 1 mM MgCl₂, and 5% glycerol; for FtsZ2-1, 25 mM glycine (pH 10.5), 50 mM KCl, 1 mM MgCl₂, and 5% glycerol. This resulted in highly enriched FtsZ1-2 or FtsZ2-1, respectively (Supplemental Figure S8).

### 3.5. *In vitro*-polymerization of Physcomitrella FtsZ1-2 and FtsZ2-1

To evaluate the activity of both FtsZ proteins, polymer formation was assessed using light-scattering assays. This method is highly effective for monitoring FtsZ filament formation in real-time, as the intensity of light scattering is directly proportional to the mass of polymers formed [71]. The light-scattering experiments were performed using the optimized purification buffers. For both proteins, a protein concentration-dependent increase in light scattering was observed upon the addition of GTP, indicating the formation of FtsZ filaments (Figure 4A and B). Interestingly, both proteins exhibited an initial lag phase following GTP addition, during which no increase in light scattering was detected. Attempts to enhance the polymerization rate of the two FtsZ proteins by altering the buffer conditions to those typically favouring polymerizations in other FtsZ proteins, such as lowering the pH or increasing the potassium concentration, resulted in protein precipitation. However, when equal concentrations of the two FtsZ proteins were combined, a significantly faster increase in light scattering was observed after GTP addition, indicating a synergistic interaction between the two proteins (Figure 4 C).

**Figure 4.**
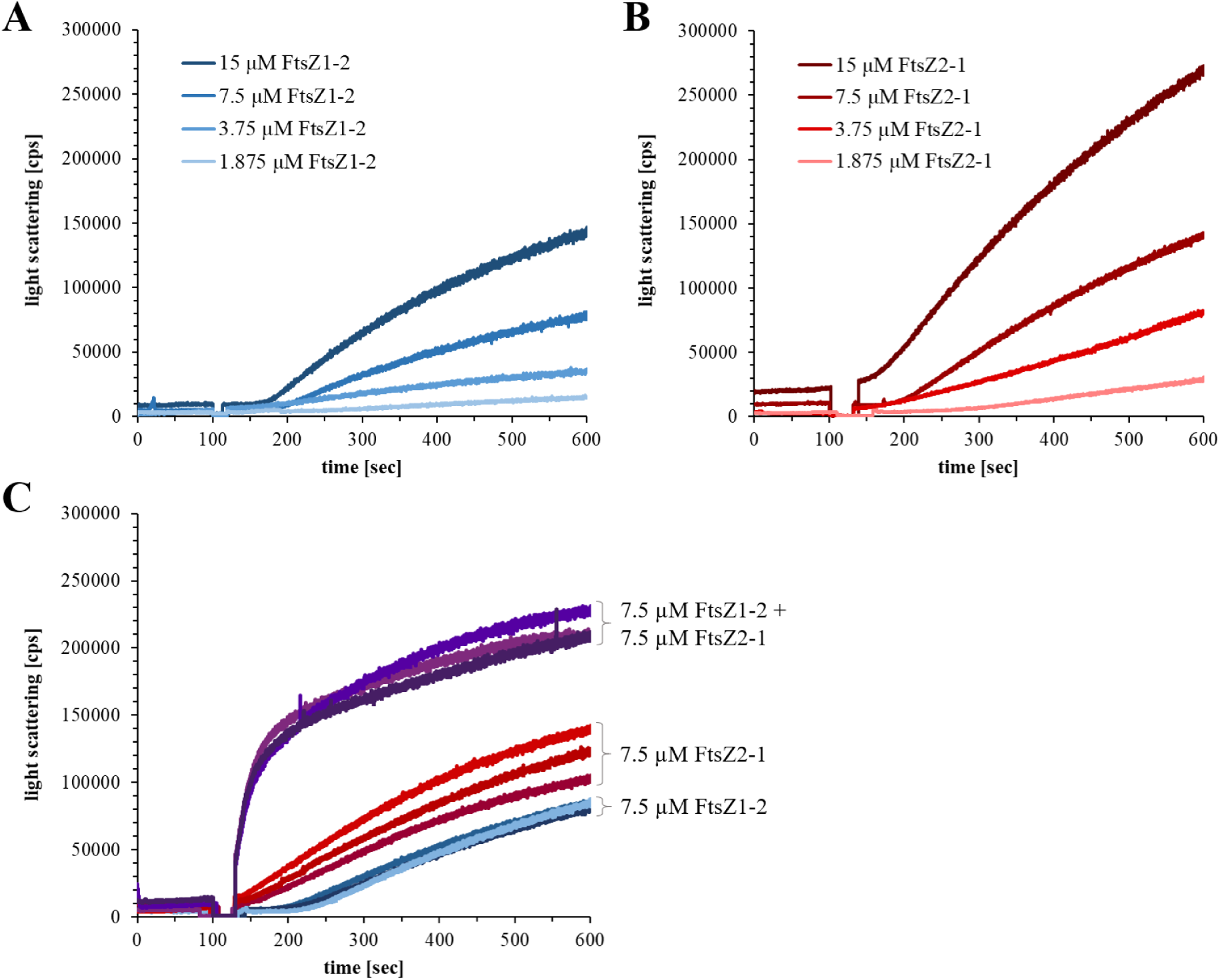
Light scattering experiments with FtsZ1-2 and FtsZ2-1, individually and in combination. Light scattering experiments were conducted at 25 °C and GTP was added after 100 sec at a final concentration of 5 mM. Individual light scattering experiments were performed using 1.875 µM, 3.5 µM, 7.5 µM or 15 µM of either of FtsZ1-2 in buffer C (**A**) or FtsZ2-1 in buffer B (**B**). The combined light scattering experiments were performed in a buffer containing 37.5 mM glycine (pH 10), 100 mM KCl, 5 mM MgCl₂, and 5% glycerol. For these experiments, 7.5 µM of each FtsZ1-2 and FtsZ2-1 were mixed and analysed in technical triplicates. Additionally, 7.5 µM of each FtsZ variant was measured separately in triplicates under the same buffer conditions (**C**). Light scattering in counts per seconds (cps).

### 3.6. Physcomitrella FtsZ proteins show GTPase activity *in vitro*

To further investigate the synergistic interaction between FtsZ1-2 and FtsZ2-1, the GTPase hydrolysis rates of the proteins were measured. Both FtsZ proteins exhibited low but significant GTPase activities. FtsZ1-2, which demonstrated a lower and slower increase in light scattering, displayed higher GTPase activity compared to FtsZ2-1. Stoichiometric mixing of both FtsZ isoforms led to a significant increase in GTPase activity, exceeding the activity observed for the individual proteins (Figure 5). This observation further supports the existence of a synergistic interaction between the two FtsZ isoforms.

**Figure 5.**
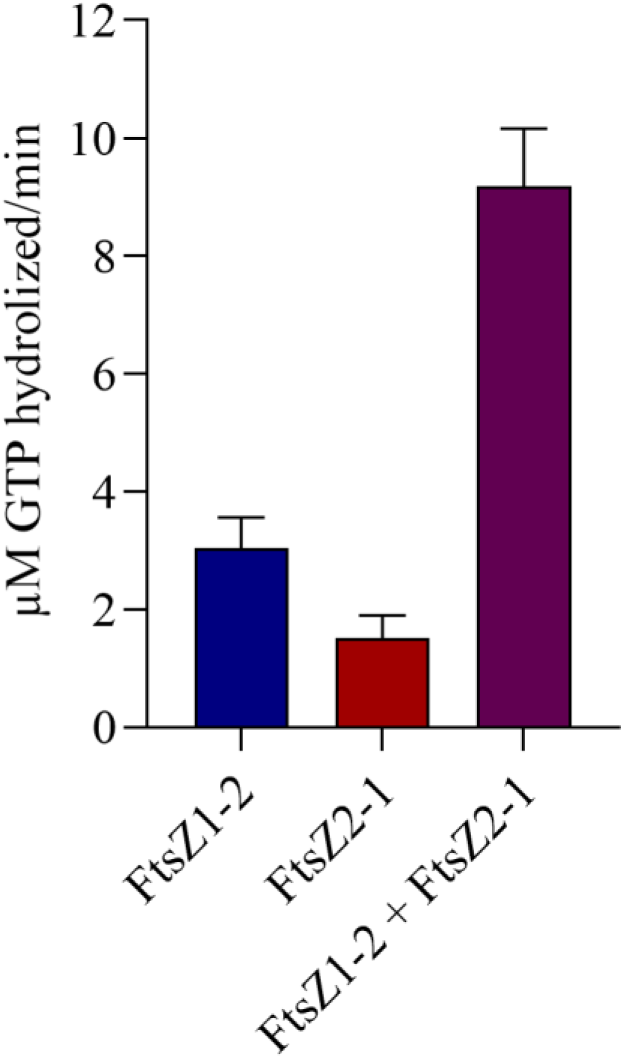
GTPase activity of Physcomitrella FtsZ1-2 and FtsZ2-1, individually and in combination. GTPase assays were performed at 25 °C using either 7.5 µM of FtsZ1-2 or 7.5 µM of FtsZ2-1 individually, or a mixture of 7.5 µM FtsZ1-2 with 7.5 µM FtsZ2-1. The assays were performed in a buffer containing 37.5 mM glycine (pH 10), 100 mM KCl, 5 mM MgCl₂, and 5% glycerol in the presence of 1 mM GTP and conducted in technical triplicates. To account for background GTP auto-hydrolysis, parallel reactions without FtsZ variants were included for each replicate. Background values were subtracted from the corresponding experimental measurements. Absorbance was measured at 620 nm after a 10-minute incubation. The amount of free phosphate (µM) released per minute was determined using a phosphate standard calibration curve. Data are presented as mean ± standard deviation from three technical replicates.

### 3.7 GTP-dependent filament formation of FtsZ1-2 and FtsZ2-1

The formation of FtsZ filaments was further investigated using negative-stain transmission electron microscopy. Under the same conditions used for the GTPase and light-scattering experiments, filaments were observed for both FtsZ1-2 and FtsZ2-1 when incubated in the presence of 2 mM GTP (Figure 6), while no filaments were detected in the absence of GTP (Supplemental Figure S9). FtsZ1-2 formed small protofilaments, which further assembled into wider filaments, adopting a straight, bundle-like conformation. In contrast, FtsZ2-1 mainly polymerized into curved filaments with straight filaments being observed only occasionally. While filament bundles formed from FtsZ1-2 were wider, those formed from FtsZ2-1 were notably thinner. When FtsZ1-2 and FtsZ2-1 were co-incubated in the presence of GTP, both curved and straight filaments were observed with occasionally single protofilaments also present. Overall the appearance of the filaments more closely resembled that of FtsZ2-1. However, it was not possible to distinguish between FtsZ1-2, FtsZ2-1, or potential hybrid FtsZ1-2/FtsZ2-1 filaments (Figure 6). In conclusion, purified Physcomitrella FtsZ1-2 and FtsZ2-1 are active and capable of forming FtsZ filaments in the presence of GTP.

**Figure 6.**
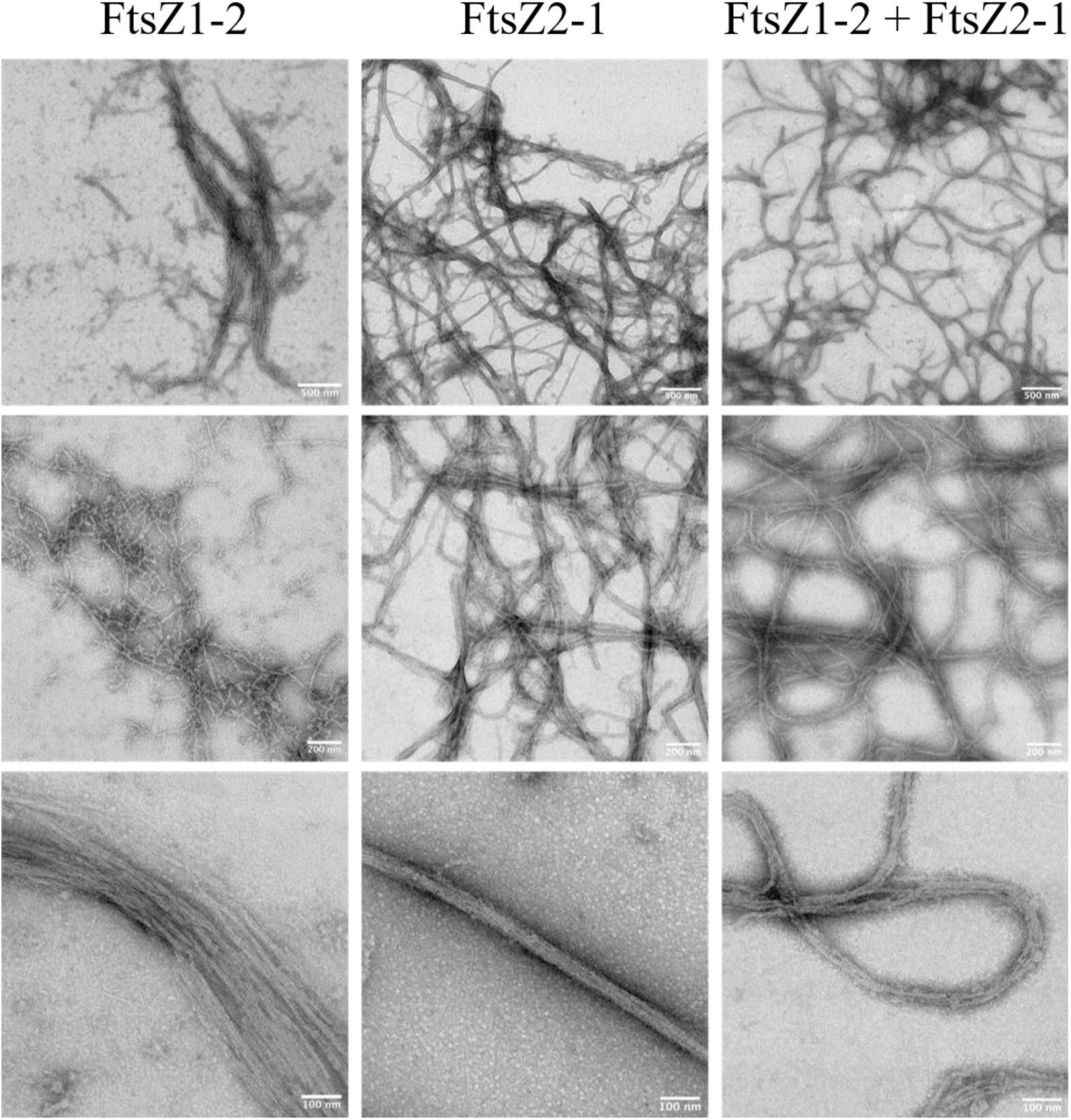
Negative stain transmission electron microscopies of Physcomitrella FtsZ filaments. 7.5 µM of FtsZ1-2 (left), 7.5 µM FtsZ2-1 (middle) and 7.5 µM FtsZ1-2 mixed with 7.5 µM FtsZ2-1 (right) were incubated at room temperature for 5 min with 2 mM GTP and then imaged by transmission electron microscopy. Representative images are shown. Scale bars as indicated. Experiments were repeated with two independent purifications with similar results.

## 4. Discussion

FtsZ proteins are GTPases and GTP binding is a prerequisite for their polymerization, and GTP hydrolysis supports filament dynamics and turnover [21, 22, 34]. The moss Physcomitrella encodes five different FtsZ proteins, suggesting neofunctionalisation of the different isoforms during evolution [30, 39, 40]. The interplay among the five FtsZ isoforms results in the formation of a filament network inside the chloroplasts, where each isoform exhibits distinct filament characteristics [52]. Two of the five isoforms (FtsZ1-2, FtsZ2-1) have relatively high expression levels in various moss tissues, indicating division-independent, stable networks within the chloroplast [53]. Knowledge about the polymer-forming properties of FtsZ1-2 and FtsZ2-1 as a GTP-driven non-equilibrium structure is of particular interest to develop sustainable material systems [73]. Here, we expressed FtsZ1-2 and FtsZ2-1 in *E. coli*. We studied the localization of fluorescence-tagged Physcomitrella FtsZ in *E. coli* cells, established a purification protocol for the recombinant FtsZ isoforms, and characterized their GTPase activity and polymerization *in vitro*. Expression of fluorescence-tagged Physcomitrella FtsZ1-2 and FtsZ2-1 in *E. coli* resulted in fluorescent foci and elongated cells. The distribution pattern of fluorescence-tagged Physcomitrella FtsZ is similar to *E. coli* FtsZ tagged with GFP [74]. Elevated levels of *E. coli* FtsZ tagged with GFP also localize in foci, and overexpression of *E. coli* FtsZ at least 10-fold above wild-type levels inhibits cell division, leading to filamentous cell shapes. In contrast, mild overexpression induces the formation of minicells [74, 75]. The foci at higher FtsZ concentrations were suggested to represent FtsZ nucleation sites [74]. In contrast, *Synechococcus elongatus* FtsZ tagged with mCerulean and Arabidopsis FtsZ1 tagged with mVenus localize to long curved filaments when expressed separately in yeast [23]. The localization of Physcomitrella FtsZ1-2 and FtsZ2-1 in *E. coli* indicates that the FtsZ function is evolutionary conserved and that Physcomitrella FtsZ interferes with cell division and morphology when overexpressed in *E. coli*.

To analyse the *in vitro*-polymerization of both Physcomitrella FtsZ isoforms individually and in combination, GTPase assays and light scattering assays were performed. GTPase and light scattering assays are FtsZ concentration dependent and are generally performed at concentrations between 1 and 15 µM. For example, concentrations ranging up to 8 µM for *Agrobacterium tumefaciens* FtsZ [63] and 12 µM for *Bacillus subtilis* FtsZ and *E. coli* FtsZ [60] have been used previously. Notably, *E. coli* FtsZ polymerizes rapidly and reaches a steady-state within only 30 seconds [72]. Indeed, both Physcomitrella FtsZ proteins exhibited concentration-dependent polymerization at concentrations higher than ∼ 2 µM. While the individual proteins exhibited a short lag phase prior to polymerization, a mixture of both proteins initiated polymerization immediately. Although strong polymerization was observed, the GTPase activity (∼0.2-0.4 GTP/min) was relatively low compared to that reported for FtsZ proteins from other organisms [60]. This may, however, be attributed to the instability of both FtsZ proteins at low and neutral pH as well as under high-salt conditions, limiting the assays to high-pH and low-salt conditions. These conditions deviate from both the natural environment and the conditions under which most FtsZ proteins exhibit optimal polymerization activity. GTP hydrolysis is essential for depolymerization, and consistent with this, no depolymerization of the FtsZ proteins was observed during the 10-minute duration of the experiments. Alternatively, our results support the idea [23] that a low GTPase activity of plant FtsZ proteins compared to bacterial FtsZ is a conserved feature. One reason might be that bacteria divide faster than chloroplasts, and thus require a higher GTPase activity and faster assembly of the Z-ring. In contrast to bacteria, chloroplasts are not able to divide on their own as chloroplast division relies on additional nuclear-encoded proteins [76]. This implies that only the initiation of chloroplast division itself might take as much time as the whole division process in bacteria. Furthermore, plant-specific components of the division machinery are involved in the rate of chloroplast division, e.g. PLASTID DIVISION (PDV) proteins. The overexpression of PDV proteins in both Arabidopsis and Physcomitrella led to an increase in chloroplast number and a decrease in chloroplast size [76]. Additionally, the rates of chloroplast division might vary depending on the developmental stage: during leaf development the level of PDV proteins decreased, leading to a decrease in the chloroplast division rate [76]. This indicates another mechanism determining the rate of chloroplast division, which might explain why chloroplasts take more time to divide, therefore not depending on a high FtsZ GTPase activity. Thus, the relatively slow GTP-dependent polymerization of the Physcomitrella FtsZ proteins at relatively high concentrations observed here is in accordance with their participation in a plastoskeleton, which persists independent of the division process.

By performing Co-IPs, we confirmed that Physcomitrella FtsZ2-1 interacts with Physcomitrella FtsZ1 proteins *in vivo*. In FRET experiments, the *in-vivo* interaction between Physcomitrella FtsZ2-1 and FtsZ1-1 was observed before [52]. Our Co-IPs were performed with an in-frame fusion of Physcomitrella FtsZ2-1 with GFP at the endogenous locus and not with over-expression lines. Due to the relatively low abundance and high similarity of these isoforms, the specific interacting FtsZ1 isoform could not be unequivocally identified based on the detected peptides. Consequently, it was decided not to follow up on the approach to estimate the ratio of FtsZ1-2 and FtsZ2-1 forming (hetero)polymers but to perform GTPase assays and polymerization assays with a mixture of Physcomitrella FtsZ1-2 and FtsZ2-1 in a 1:1 ratio. This mixing resulted in a significantly enhanced overall GTPase activity, suggesting that there is a synergistic interaction of the two FtsZ isoforms. This result is different from findings regarding Arabidopsis FtsZ proteins [34], where no synergistic effects were observed.

We were able to visualize GTP-dependent filament formation for both Physcomitrella FtsZ isoforms using negative stain TEM, revealing distinct structural characteristics. For FtsZ1-2 small protofilaments and larger filaments with a bundle-like architecture were observed, indicating that FtsZ1-2 primarily polymerizes into small protofilaments, which subsequently assemble into straight filament bundles. In contrast, FtsZ2-1 predominantly formed curved filaments, with straight filaments being less frequent. No protofilaments were observed for this isoform. Additionally, a higher abundance of FtsZ2-1 filaments was observed in TEM images compared to FtsZ1-2. This is consistent with the light-scattering data, which demonstrated that FtsZ2-1 achieved a higher scattering intensity (measured in counts per second) than FtsZ1-2. The structural differences extended to filament thickness, where FtsZ1-2 bundles were notably broader than those of FtsZ2-1. When both isoforms were co-incubated in the presence of GTP, the resulting filaments displayed characteristics of both straight and curved morphologies, along with occasional single protofilaments. Interestingly, the overall filament appearance was more similar to FtsZ2-1. However, it was not possible to definitively attribute the observed filaments to FtsZ1-2, FtsZ2-1, or hybrid assemblies. Differences in the polymer-forming abilities of plant FtsZ proteins have been previously reported. Arabidopsis FtsZ2 forms filaments similar to those of Physcomitrella FtsZ1-2 as observed by TEM, whereas Arabidopsis FtsZ1 does not independently assemble into filaments under the tested conditions [34]. In Physcomitrella, FtsZ2-1 is directly involved in plastid division [8, 41] leading to enlarged plastids in basal chloronema cells of the Δ*ftsZ2-1* mutant [41]. Furthermore, chloroplasts of Δ*ftsZ2-1* cells show complete loss of chloroplast integrity [53]. In contrast, chloroplasts in the chloronema cells of the Δ*ftsZ1-2* mutant did not show morphological differences compared to WT cells [53]. This indicates that FtsZ2-1 is essential for maintaining chloroplast shape and integrity, while FtsZ1-2 might be less important. While filaments of FtsZ2-1 are exclusively formed within the chloroplast, FtsZ1-2 filaments apparently need to be longer, as they form long, extra-plastidic extensions to connect multiple chloroplasts *via* stromules [43, 53]. In general, it is reasonable to assume that differences in filament morphology are due to the underlying differences in GTPase activity and polymerization dynamics.

### Conclusions

We observed GTPase activity and *in-vitro* polymerization of Physcomitrella FtsZ1-2 and FsZ2-1. We confirmed the polymer formation of both FtsZ isoforms using TEM and show that Physcomitrella FtsZ1 isoforms were identified as significant interactors of Physcomitrella FtsZ2-1 *in vivo*. We present the heterologous expression of eGFP-tagged Physcomitrella FtsZ1-2 and mKO2-tagged FtsZ2-1 in *E. coli*. The observation that the over-expression of both Physcomitrella FtsZ isoforms in *E. coli* leads to elongated bacterial cells confirms that the FtsZ function for bacterial and organelle division is evolutionary conserved. Furthermore, our results provide insights into the polymer-forming properties of Physcomitrella FtsZ1-2 and FtsZ2-1. Based on the mutant phenotypes [42, 53] the localization of Physcomitrella FtsZ in *E. coli* (this study), the *in vivo* ([53] and this study) and our *in-vitro* studies, we postulate different roles for FtsZ1-2 and FtsZ2-1 regarding the assembly of the plastoskeleton. These diverse functions of FtsZ1-2 and FtsZ2-1, as well as their interaction, impact the formation of the Physcomitrella plastoskeleton, abidingly shaping chloroplast morphology.

## 5. Methods

### 5.1. Design of the expression vectors FtsZ1-2 and FtsZ2-1

The coding sequences (CDS) for *Pp*FtsZ1-2 (Pp3c19_2490) and *Pp*FtsZ2-1 (Pp3c11_17860) according to the Physcomitrella genome [78] were optimized for bacterial expression. Changes in the CDS were made accordingly to a codon usage table derived from the analysis of 16,000 *E. coli* genes using a custom Perl script and the GC content was concurrently increased (script written by Dr. Oguz Top). The putative N-terminal transit peptides for *Pp*FtsZ1-2 (Pp3c19_2490) and *Pp*FtsZ2-1 (Pp3c11_17860) were determined using the ChloroP 1.1 Prediction Server (http://www.cbs.dtu.dk/services/ChloroP/). The codon-optimized sequences lacking the predicted N-terminal transit peptides (34 and 32 amino acids, respectively, Table 1) were synthesized by Eurofins Genomics Germany GmbH, Ebersberg, Germany. For cloning of the final construct of FtsZ2-1, the primers were designed starting at the codon for amino acid 33. The fragments for FtsZ1-2 and FtsZ2-1 were amplified via PCR using the respective primers (Supplemental Tables S3 and S4). The final expression vectors were subcloned into the pLATE11 bacterial expression vector via Ligation Independent Cloning (LIC, aLICator LIC Cloning and Expression Kit 1, untagged; Thermo Fisher Scientific, Waltham, USA) following the manufacturer’s instructions. The pLATE11 expression vector allows high levels of target protein expression. The 8 x His-tag was integrated into the primer and added either to the N-terminus or the C-terminus of the CDS.

### 5.2. Design of the expression vectors pLATE_eGFP_His, pLATE_eGFP_FtsZ1-2_His, pLATE_mKO2_His, pLATE_mKO2_FtsZ2-1_His

To generate the expression vectors pLATE_eGFP_His, pLATE_eGFP_FtsZ1-2_His, pLATE_mKO2_His, pLATE_mKO2_FtsZ2-1_His, the CDS of FtsZ1-2, FtsZ2-1, eGFP and mKOrange2 (mKO2) were amplified via PCR using the respective primers (Supplemental Table S3 and S4). The mKO2 template was kindly provided by Dr. Roland Nitschke. The inserts were then assembled via Gibson cloning [80] and subcloned into the pJET1.2 vector (CloneJET PCR cloning kit, Thermo Fisher Scientific) following the manufacturer’s instructions. After identification of the recombinant clones, the correct construct was used as a template to be amplified using the respective primers for LIC cloning (Supplemental Table S3). The PCR fragments were subcloned into the pLATE11 expression vector via Ligation Independent Cloning following the manufacturer’s instructions. To create the construct containing both fluorescence-tagged FtsZ isoforms, the pLATE_eGFP_FtsZ1-2_His vector was linearized using Nde I. Subsequently, the mKO2-FtsZ2-1 expression cassette, amplified from the pLATE_mKO2_FtsZ2-1_His vector, was inserted into the corresponding site by Gibson cloning.

### 5.3. Design of the expression vectors His_6__SUMO_FtsZ1-2 and His_6__SUMO_FtsZ2-1

The CDS of FtsZ1-2 and FtsZ2-1 lacking the N-terminal transit peptides (86 and 42 amino acids, respectively) were amplified via PCR (Supplemental Table S4) and subcloned into the linearized pSVA13429 vector [76] via Gibson cloning [80]. The final vectors were sequenced with SUMO_fwd and T7_rev (Supplemental Table S4).

### 5.4. Transformation of E. coli cells

BL21 Star™ (DE3) One Shot® (Invitrogen, Thermo Fisher Scientific) or NiCo (Invitrogen, Thermo Fisher Scientific) cells were transformed with vectors listed in Table 2 according to the manufacturer’s instructions.

**Table 2.** Plasmids used in this study.

| Plasmid | Relevant characteristics |
| --- | --- |
| pLATE_FtsZ1-2_His<sub>8</sub> | Overproduction FtsZ1-2 with C-terminal His<sub>8</sub> |
| pLATE_FtsZ2-1_His<sub>8</sub> | Overproduction FtsZ2-1 with C-terminal His<sub>8</sub> |
| pLATE_eGFP_His<sub>8</sub> | Overproduction eGFP with His<sub>8</sub> |
| pLATE_eGFP_FtsZ1-2_His<sub>8</sub> | Overproduction FtsZ1-2 with N-terminal eGFP and C-terminal His<sub>8</sub> |
| pLATE_mKOrange2_His<sub>8</sub> | Overproduction mKOrange2 with His<sub>8</sub> |
| pLATE_mKOrange2_FtsZ2-1_His<sub>8</sub> | Overproduction FtsZ2-1 with N-terminal mKOrange2 and C-terminal His<sub>8</sub> |
| pLATE_eGFP_FtsZ1-2_His<sub>8</sub>_mKOrange2_FtsZ2-1_His<sub>8</sub> | Overproduction FtsZ1-2 with N-terminal eGFP and C-terminal His<sub>8</sub> and overproduction FtsZ2-1 with N-terminal mKOrange2 and C-terminal His<sub>8</sub> |
| His<sub>6</sub>_SUMO_FtsZ1-2 | Overproduction FtsZ1-2 with N-terminal His<sub>6</sub> and SUMO-tag |
| His<sub>6</sub>_SUMO_FtsZ2-1 | Overproduction FtsZ2-1 with N-terminal His<sub>6</sub> and SUMO-tag |
| pLATE31-Cm | Overproduction CAT |

### 5.5. Generation of a FtsZ2-1-eGFP fusion line

The coding sequence of eGFP combined with a flexible 18 bp linker [82] fused FtsZ2-1 at the endogenous locus via homologous recombination as described [81]. Homologous flanks for the integration were chosen to remove the endogenous stop codon. Restriction sites for Esp3I to release the linearized transfection construct from the final vector backbone were included at the 5’ and the 3’ end of the knock-in construct. All necessary parts were amplified with Phusion polymerase (Thermo Fisher Scientific) and assembled into the pJet1.2 vector backbone (Thermo Fisher Scientific) via triple template PCR as described [82]. PCRs were performed with the primers listed in Supplemental Table S1. The plasmid linearized *via* Alw26I digestion as well as a plasmid for co-transfection containing a neomycin resistance cassette [83] were purified via ethanol precipitation and transfection and selection was performed as described [64, 84]. Plants surviving the selection procedure were screened by PCR (Phire Polymerase, Thermo Fisher Scientific) for targeted integration of the knock-in construct at the desired locus as described [81]. Sequences of primers used for cloning and screening are listed in Supplemental Table S3. Physcomitrella WT as well as the employed eGFP fusion line #516 is available from the International Moss Stock Center (IMSC, www.moss-stock-center.org) with accessions 40095 and 40960, respectively. The plasmid containing the knock-in construct is also available from the IMSC under accession P1779.

### 5.6. Co-immunoprecipitation

Co-immunoprecipitations of the GFP-tagged FtsZ2-1 line was performed in biological triplicates using GFP-Trap Magnetic Particles M-270 (ChromoTek, Planegg-Martinsried, Germany). In brief, 300 mg homogenized protonema of the bait line (GFP-tagged FtsZ2-1, line #513) and WT were dissolved in 2 mL ice-cold extraction buffer (25 mM HEPES-KOH, pH 7.5, 2 mM EDTA, 100 mM NaCl, 200 nM DTT, 0.5 % Triton X-100, 1 % plant protease inhibitor cocktail (PPI, P9599, Sigma Aldrich, St. Louis, USA). All further steps were performed as described [81]. MS analyses were performed on a Q-Exactive Plus instrument (Thermo Fisher Scientific) coupled to an UltiMate 3000 RSLCnano system (Dionex, Thermo Fisher Scientific) as described [85]. A database search of the test Co-IPs was performed with Mascot Server V2.7.0 (Matrix Science). Results were subsequently loaded into the Scaffold^TM^ 5 software (V5.0.1, Proteome Software) and proteins were accepted at an FDR = 1 and peptides at an FDR = 0.5. Database search and label free quantitation of the quantitative CoIP were performed as described [81] using *MaxQuant* (2.1.4.0; Cox and Mann, 2008). The database contained all Physcomitrella V3.3 protein models [78] as well as a fusion sequence of FtsZ2.1 (Pp3c11_17860) and eGFP Data analysis was performed in Perseus (V2.0.10.0 [83]). Proteins with at least two LFQ values in at least one group (FtsZ or WT) were used and missing values were imputed from a normal distribution with a down-shift of 1.8 and distribution width of 0.5. True interaction partners were accepted at an FDR of 0.01 % and a p-value < 0.01. The resulting table containing significantly interacting proteins is available from Supplemental Table S2.

### 5.7. Preparation of transformed BL21 Star™ (DE3) cells for confocal microscopy

For confocal microscopy, 3 mL of LB medium containing 50 mg/mL ampicillin (amp) and 1 % glucose was inoculated with a colony of respective transformed BL21 Star™ (DE3) cells, picked from a LB/amp agarose plate. The inoculated culture was then incubated overnight at 37° C while shaking. On the subsequent day, 200 µL of this overnight culture was transferred to 2.8 mL of LB/amp and further incubated for 3 hours at 37° C. Subsequently, 0.5 mM isopropyl ß-D-1-thiogalactopyranoside (IPTG) was added, and the culture was incubated for an additional 45 minutes. For confocal microscopy, 1-5 µL of the respective bacterial solution was transferred to a microscopy glass slide. Then the samples were embedded using ProLong™ Glass Antifade Mountant (Thermo Fisher Scientific), covered with a cover slide, and stored in darkness for 24 hours prior to confocal microscopy.

### 5.8. Confocal microscopy of transformed BL21 Star™ (DE3) cells

All images were taken with a Zeiss LSM880 NLO microscope (Carl Zeiss Microscopy GmbH, Jena, Germany) using Plan-Apochromat 40x/1.4 Oil DIC(UV)VIS-IR objective with a zoom factor of 2. For the excitation the laser was applied at 488 nm (eGFP) or 561 nm (mKO2) with an intensity of 2 % or 5 %, respectively. For eGFP images the pinhole was adjusted to 100.5 AU (12.11 µm), and for mKO2 the pinhole was set to 115.4 AU (18.54 µM), while no emission filter was used. The detection range was set to 500-550 nm for the eGFP fluorescence and 560-600 nm for the mKO2 fluorescence.

### 5.9. Overexpression and purification of His_6__SUMO_FtsZ1-2 and His_6__SUMO_FtsZ2-1

NiCo cells (Invitrogen, Thermo Fisher Scientific) were transformed with the plasmids His₆_SUMO_FtsZ1-2 and His₆_SUMO_FtsZ2-1, then directly transferred to 200 mL of LB medium containing 50 mg/L kanamycin and 0.025% glucose. Cultures were grown overnight at 37 °C and subsequently used to inoculate 1 litre of a simplified autoinduction medium (based on [70]) in a 5 L baffled flask, to an OD₆₀₀ of 0.025. The medium contained 100 mg/L kanamycin, 6 g/L Na₂HPO₄, 3 g/L KH₂PO₄, 10 g/L tryptone, 5 g/L yeast extract, and 5 g/L NaCl. Glucose, lactose, and glycerol were autoclaved separately as a 20× stock and added to achieve final concentrations of 3 g/L glucose, 8 g/L lactose, and 11 g/L glycerol. The culture was grown at 37 °C until reaching an OD₆₀₀ of 1.0, after which the temperature was lowered to 22 °C, and growth continued overnight. The next day, OD₆₀₀ was monitored hourly until it remained constant for 2 hours. Cells were harvested by centrifugation at 7,500 rpm (SS-34 rotor) at 4 °C, washed with either buffer A (50 mM glycine, pH 9.5, 50 mM KCl, 1 mM MgCl₂, 5% glycerol) containing 20 mM imidazole for FtsZ1-2, or buffer B (25 mM glycine, pH 9.5, 150 mM KCl, 1 mM MgCl₂, 5% glycerol) containing 20 mM imidazole for FtsZ2-1, and centrifuged again. The supernatant was removed, and the cell pellets were flash-frozen in liquid nitrogen and stored at −80 °C. Frozen pellets were thawed on ice and resuspended in their respective buffers (A or B) containing 20 mM imidazole. All subsequent steps, including purification, were performed on ice or at 4 °C. Cells were lysed by passing them through a French press three times, followed by centrifugation at 7,500 rpm (SS-34 rotor) and then at 45,000 rpm (Ti-60 rotor). Purification was performed using an ÄKTA Purifier system (Cytiva). The supernatant was loaded onto a 5 mL HisTrap FF column (Cytiva) equilibrated with buffer A or B containing 20 mM imidazole at a flow rate of 0.5 mL/min. The column was washed with the same buffer at 2 mL/min until the OD₂₈₀ stabilized. Proteins were eluted with buffer A or B containing 250 mM imidazole at 0.5 mL/min, and fractions containing protein were collected and pooled. The His₆-SUMO tag was removed by cleavage with His-Ulp1 protease (80 ng/mL) for 6 hours at 4 °C in the presence of 2 mM DTT, 0.1% ND-40, and 500 µM GDP. SUMO protease was produced in-house [79, 86]. Proteins were concentrated using Pierce™ Protein Concentrators PES (10 kDa MWCO, Thermo Fisher Scientific). The concentrated proteins were applied to a HiLoad® 16/600 Superdex® 200 pg (Cytiva) size-exclusion column equilibrated with buffer C (50 mM glycine, pH 10.5, 50 mM KCl, 1 mM MgCl₂, 5% glycerol) for FtsZ1-2 or buffer B for FtsZ2-1. Protein concentrations in fractions containing FtsZ1-2 and FtsZ2-1 were determined using the Pierce BCA Protein Assay Kit (Thermo Fisher Scientific) according to the manufacturer’s instructions.

### 5.10. GTP hydrolysis assay

The GTPase activity was assessed using the malachite green assay. Experiments were conducted in a buffer containing 37.5 mM glycine (pH 10), 100 mM KCl, 5 mM MgCl₂, and 5% glycerol, with a total reaction volume of 50 µL. At 25 °C, released phosphate increased linearly for at least 15 minutes and therefore the assay was performed at 25 °C for 10 minutes. The reaction was initiated by adding GTP to a final concentration of 1 mM and stopped by the addition of EDTA to a final concentration of 10 mM. A 20 µL aliquot of the reaction mixture was incubated with 80 µL of malachite green working reagent. Control reactions lacking either GTP or protein were performed under identical conditions. Absorbance at 620 nm was measured using a plate reader (CLARIOstar® Plus, BMG LABTECH GmbH, Ortenberg, Germany), and the amount of free phosphate in each sample was calculated using a phosphate standard calibration curve.

### 5.11. Light Scattering Assays

Experiments were conducted using a temperature-controlled FluoroMax-4 spectrofluorometer (Horiba Scientific) at 25 °C. The excitation and emission wavelengths were set to 350 nm, with an excitation bandwidth of 1 nm and an emission bandwidth of 0.5 nm. Filament formation was initiated by the addition of GTP to a final concentration of 5 mM. FtsZ1-2 and FtsZ2-1 were measured individually in buffers C and B in the presence of 5 mM MgCl₂, and for the experiments were FtsZ1-2 and FtsZ2-1 were mixed in a buffer D containing 37.5 mM glycine (pH 10), 100 mM KCl, 5 mM MgCl₂, and 5% glycerol.

### 5.12. Transmission electron microscopy of FtsZ filaments

For polymerization studies, FtsZ proteins (7.5 µM FtsZ1-2, 7.5 µM FtsZ2-1, or 7.5 µM each of FtsZ1-2 and FtsZ2-1) were incubated at 25 °C for 10 minutes with or without 5 mM GTP in a buffer containing 37.5 mM glycine (pH 10), 100 mM KCl, 5 mM MgCl₂, and 5% glycerol, in a total volume of 60 µL. A 5 µL aliquot of each sample was immediately applied to freshly glow-discharged carbon/Formvar-coated copper grids (300 mesh; Plano GmbH) and incubated for 30 seconds. Excess liquid was blotted away, and the grids were washed three times with ddH₂O, followed by negative staining with 2% uranyl acetate. Images were acquired using a Hitachi HT7800 transmission electron microscope operated at 100 kV and equipped with an EMSIS Xarosa 20-megapixel CMOS camera.

## 7. Acknowledgements

This project was funded by the German Research Foundation DFG under Germany’s Excellence Strategy (*liv*MatS - EXC-2193/1, Project ID 390951807, and CIBSS – EXC-2189, Project ID 390939984) to R.R. S.W.L.M. was supported by a state-funded completion scholarship of the Landesgraduiertenförderung LGFG. M.J. was supported by a grant to Sonja-Verena Albers by the Collaborative Research Centre SFB1381 funded by the DFG-Project-ID 403222702-SFB 1381. The TEM (Hitachi HT7800) was funded by the DFG grant (project number 426849454) and is operated by the University of Freiburg, Faculty of Biology, as a partner unit within the Microscopy and Image Analysis Platform (MIAP) and the Life Imaging Center (LIC), Freiburg.

We thank Oğuz Top for codon optimizing the CDS of Physcomitrella FtsZ1-2 and FtsZ2-1 for bacterial expression. We also thank the Life Imaging Center (LIC) of the University of Freiburg for help with confocal microscopy. We thank Sonja-Verena Albers for providing the equipment for purification of Physcomitrella FtsZ proteins and her overall support of this project, and Anne Katrin Prowse for language editing.

## 8. Author contributions

S.W.L.M. designed the research, performed the experiments, analysed the data, and wrote the manuscript. M.J. performed transmission electron microscopy. L.L.B. performed confocal microscopy, designed figures and helped writing the manuscript. S.N.W.H. performed quantitative Co-IPs, analysis of MS data and helped writing the manuscript. B.Ö. generated the FtsZ2-1-eGFP fusion line. E.L.D. helped designing research and writing the manuscript. C.v.d.D. designed and performed experiments, analysed data and helped writing the manuscript. R.R. designed and supervised the research, acquired funding, and wrote the manuscript. All authors read and approved the final version of the manuscript.

## Data availability

All relevant data is contained in the manuscript and in the supplementary materials. Plant material and plasmids are available from the International Moss Stock Center (IMSC, www.moss-stock-center.org) as described in the text.

## Competing Interests Statement

All authors declare no conflict of interest.

## Supplementary Information

**Table S1.**
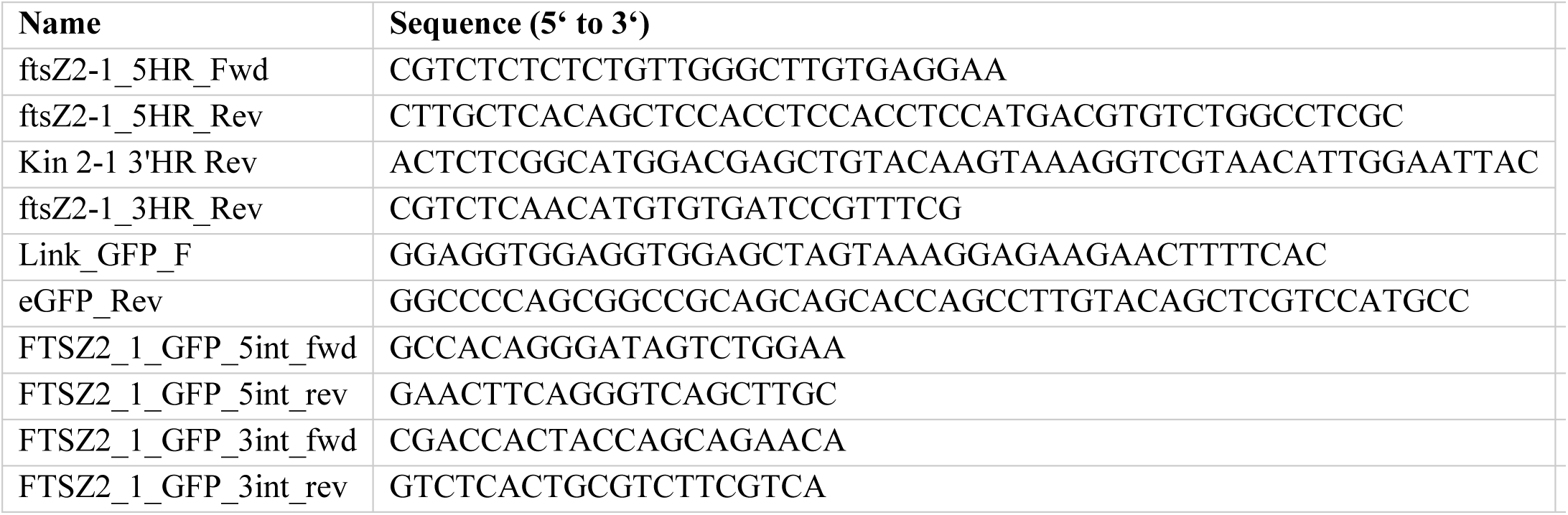
Oligonucleotides and the respective template accessions or templates used for the generation of the FtsZ2-1-eGFP fusion construct.

**Table S2 List of proteins significantly interacting with Physcomitrella FtsZ2-1 as determined by Co-IP and subsequent MS analysis** The list can be downloaded from Zenodo (https://zenodo.org/records/10648779).

**Table S3.**
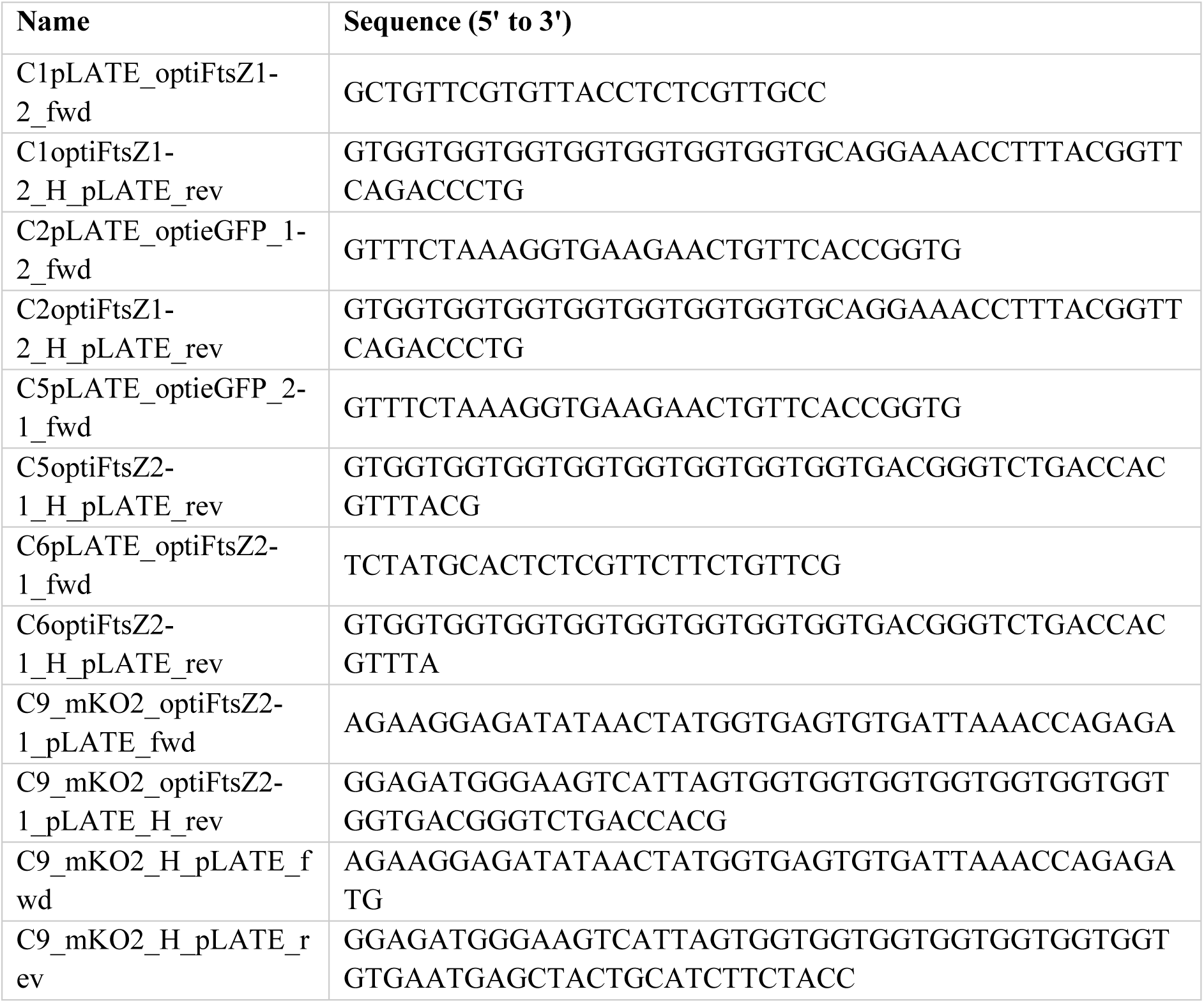
Oligonucleotides used for Ligation Independent Cloning in this work. Due to the cloning strategy, all forward oligonucleotides (fwd) include the sequence 5’ AGAAGGAGATATAACTATG 3’ at their 5’ end; all reverse oligonucleotides (rev) include the sequence 5’ GGAGATGGGAAGTCATTA 3’ at their 5’ end.

**Table S4.**
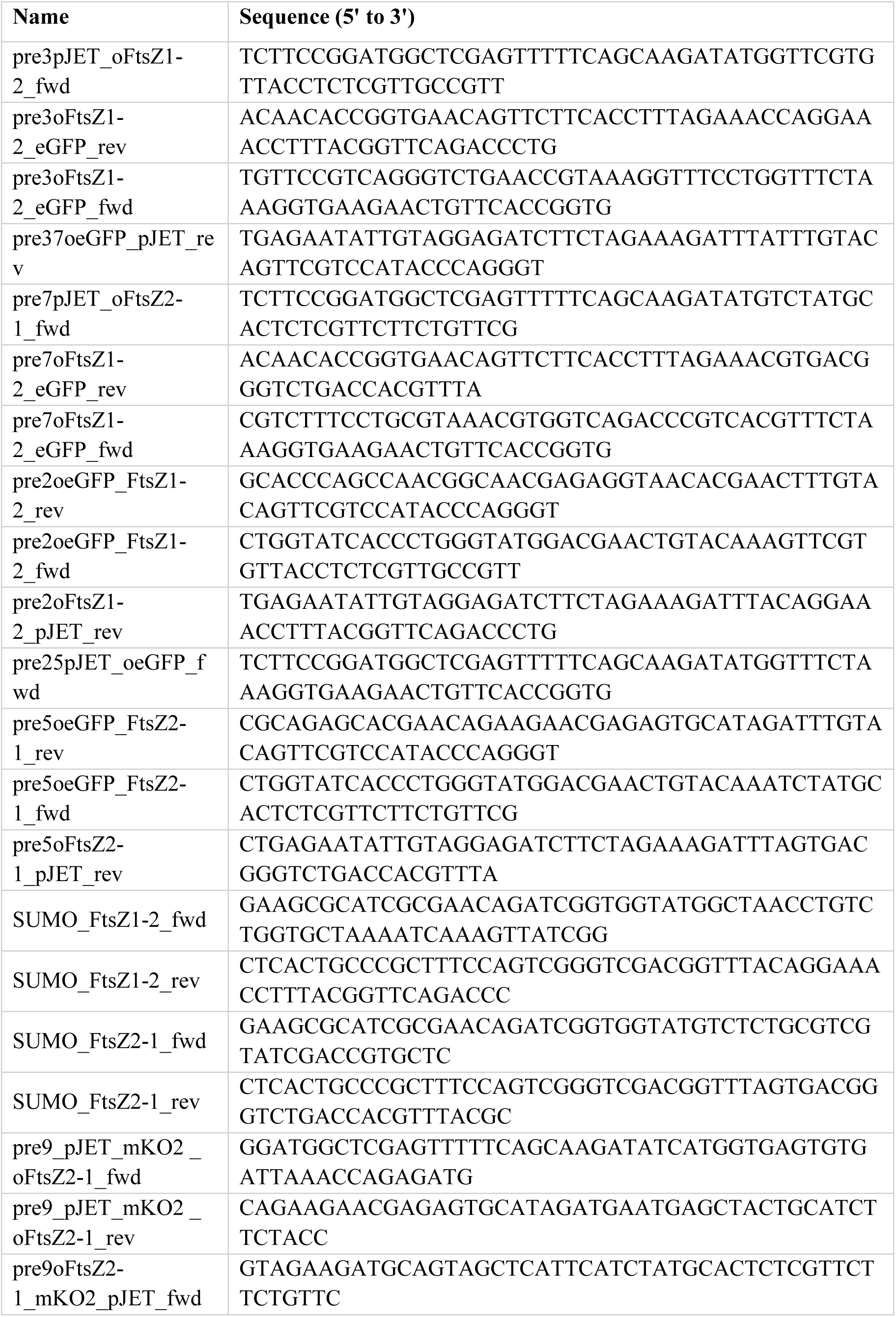
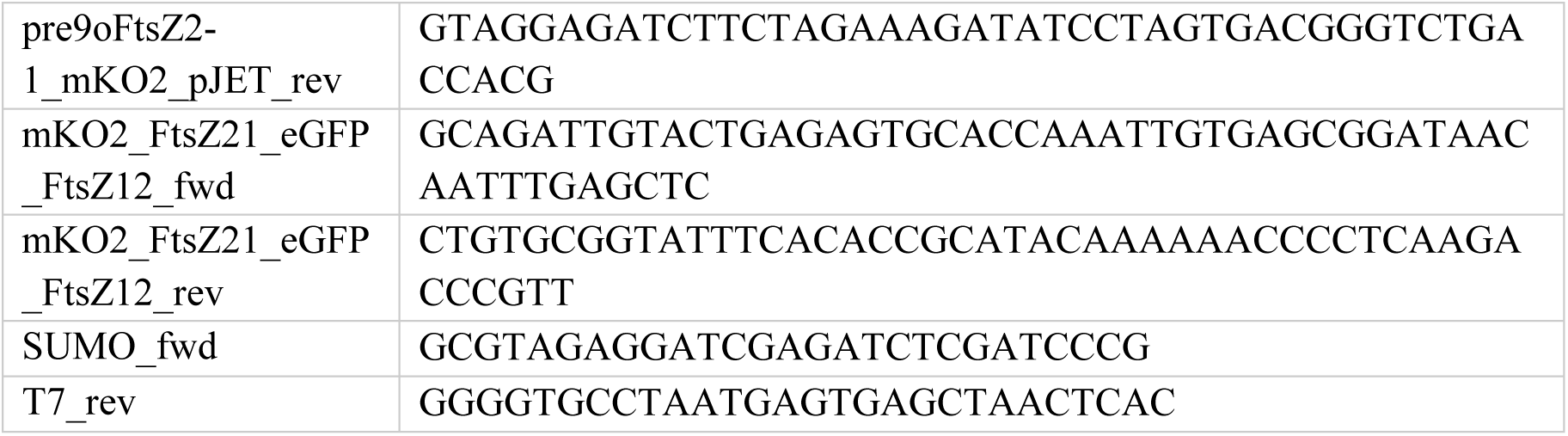
Oligonucleotides used for Gibson cloning and sequencing in this work.

**Figure S1.**
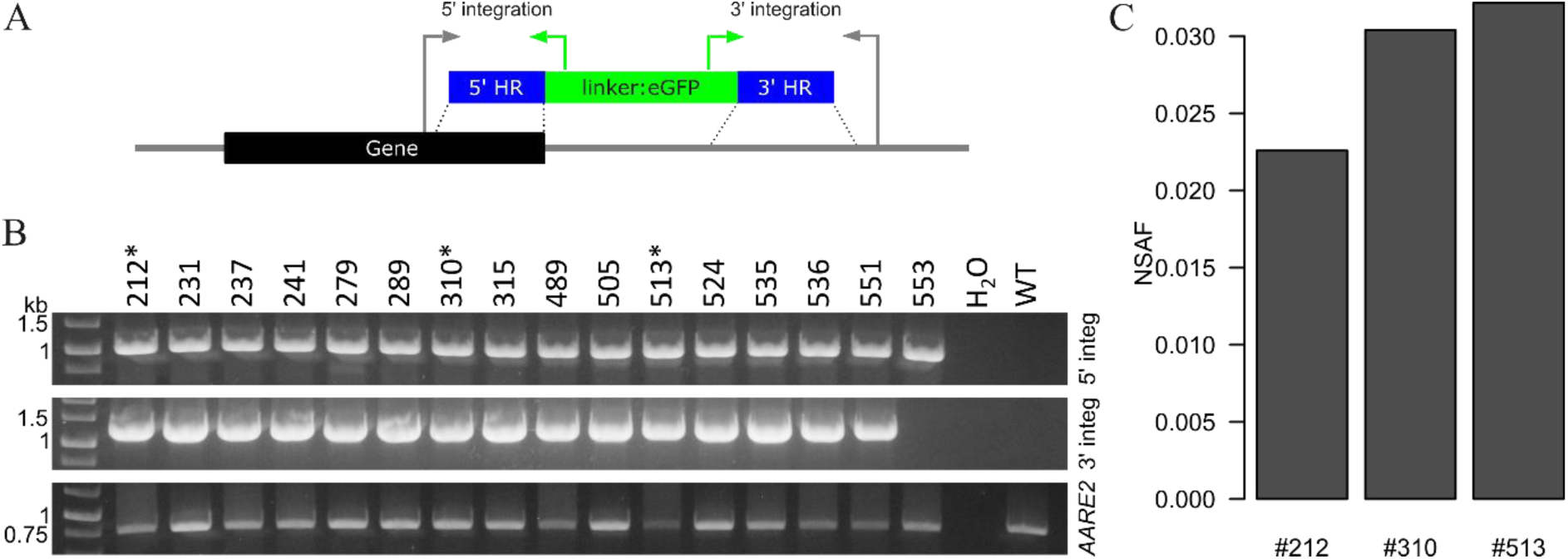
Identification of FtsZ2-1-GFP fusion lines *via* PCR and Co-immunoprecipitation (Co-IP) Primers were designed to represent correct positioning of the knock-in construct at the desired genomic locus (Pp3c11_17860V3). (**A**) Scheme representing the positioning of the knock-in construct and the primers used for the screening PCR. (**B**) Results of the screening PCR. Expected amplicon sizes: 5’ integration: 1064 bp; 3’ integration: 1302 bp; AARE2 (Pp3c12_21080V3): 768 bp. Stars (*) indicate candidate lines that were selected for a first test Co-IP. Uncropped gel images are available from Supplemental Figure S10. (**C**) Results from the first test Co-IP with selected FtsZ2-1:eGFP fusion lines. Relative quantitative share of FtsZ2.1:eGFP in the analysed sample is represented by normalized spectral abundance factors (NSAF, [87]). A full spectrum report is available from Supplemental Table S2.

**Figure S2.**
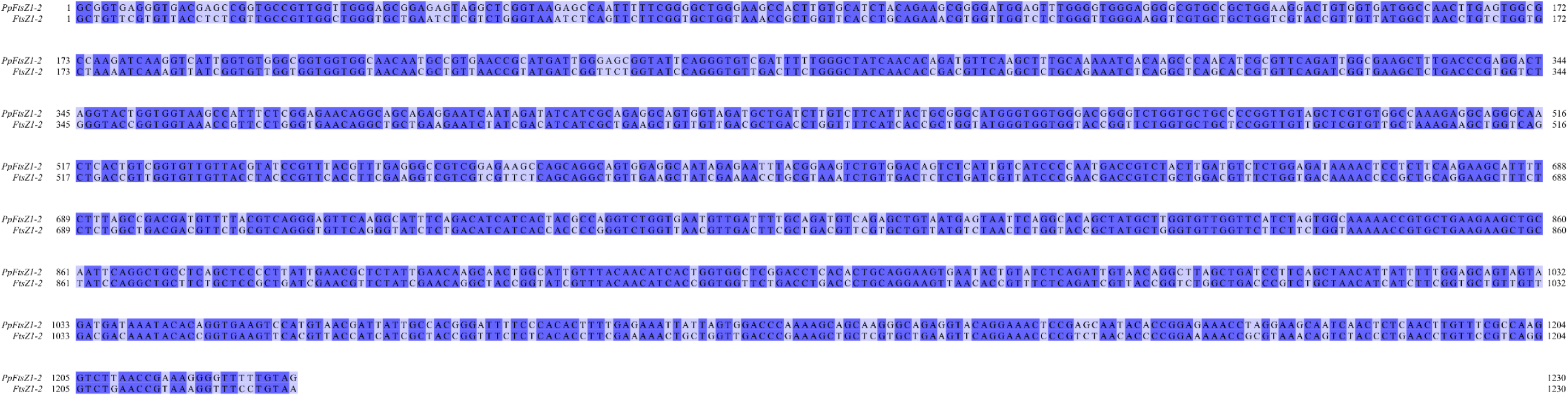
Pairwise sequence alignment of Physcomitrella FtsZ1-2 (PpFtsZ1-2) and the codon optimized FtsZ1-2 (FtsZ1-2) for codon usage in *E. coli*. Alignment was performed with Jalview (Version 2.11.2.2).

**Figure S3.**
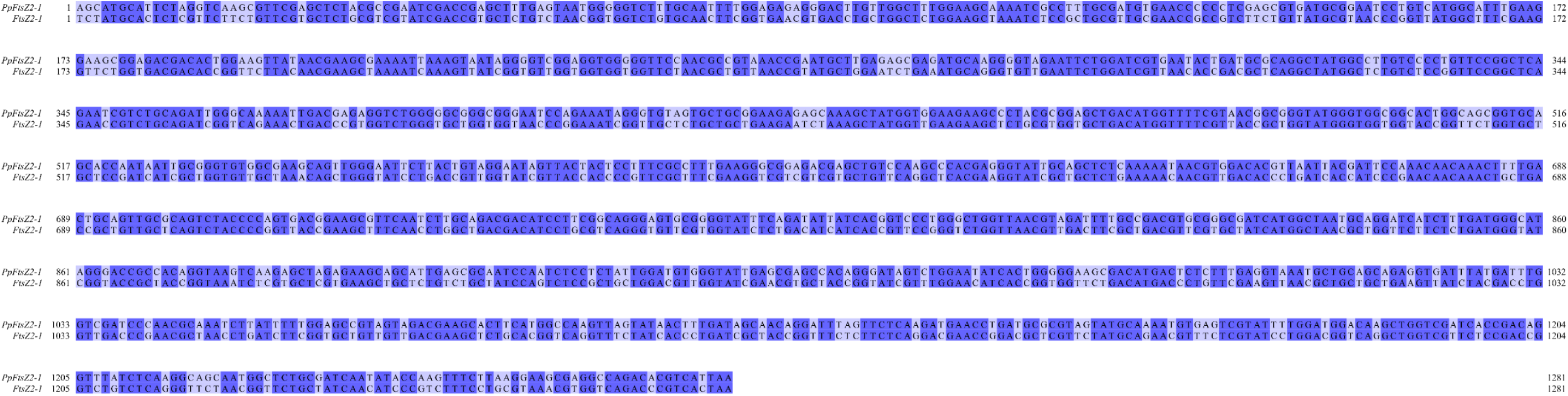
Pairwise sequence alignment of Physcomitrella FtsZ2-1 (PpFtsZ2-1) and the codon optimized FtsZ2-1 (FtsZ2-1) for codon usage in *E. coli*. Alignment was performed with Jalview (Version 2.11.2.2).

**Figure S4.**
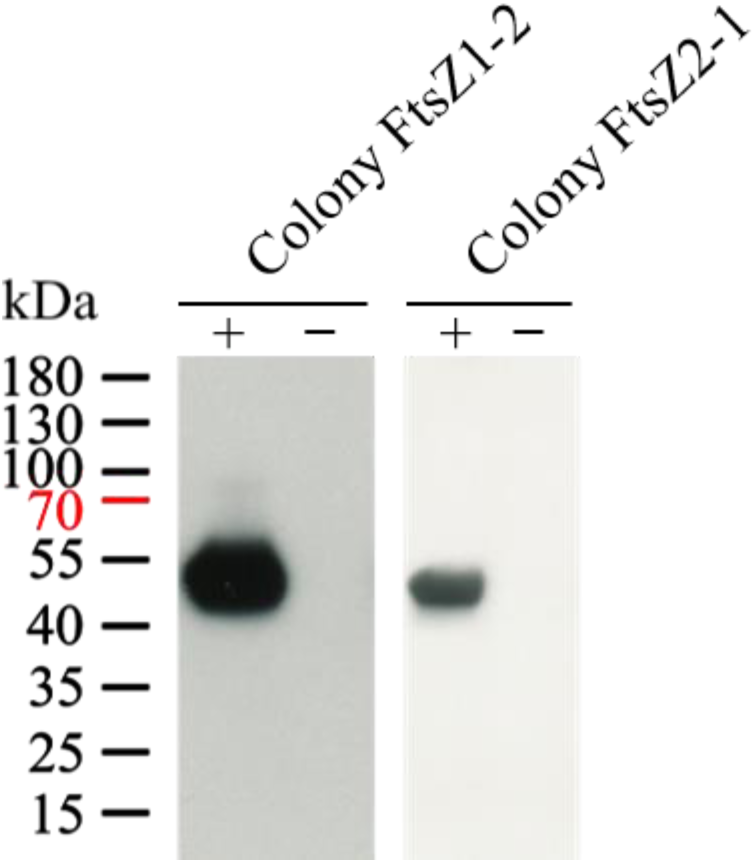
Immunoblot detections of protein extracts from BL21 Star™ (DE3) cells transformed with the pLATE11 vector for the expression of Physcomitrella FtsZ1-2 or FtsZ2-1. Total protein extracts from BL21 Star™ (DE3) cells harbouring the respective expression constructs for the Physcomitrella FtsZ isoforms were separated on a 10% SDS-PAGE gel (Bio-Rad) and subsequently transferred to a polyvinylidene fluoride (PVDF) membrane (Cytiva). Immunodetection was performed using an anti-His antibody (anti-6X His tag® antibody, ab18184, Abcam; 1:1,000) and an HRP-conjugated anti-mouse secondary antibody (NA931, Cytiva; 1:50,000). Immunoblot signals were observed at the expected molecular weights, corresponding to recombinant Physcomitrella FtsZ1-2 (∼44 kDa) and FtsZ2-1 (∼45 kDa). The ‘+’ and ‘–’ symbols indicate whether the cells were induced or not induced with IPTG. Protein ladder: PageRuler™ Prestained Protein Ladder (Thermo Fisher Scientific).

**Figure S5.**
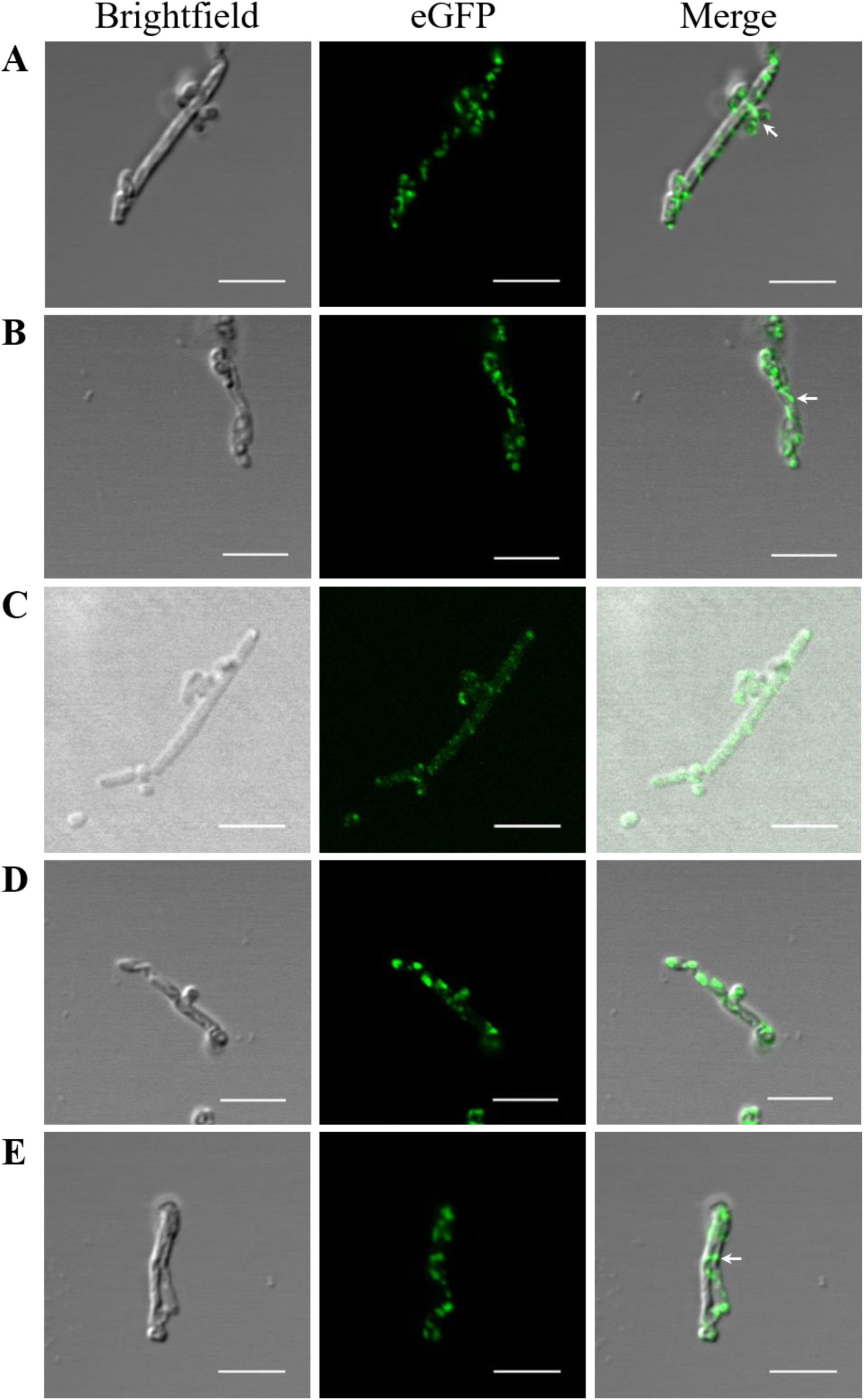
Confocal microscopy of eGFP-tagged Physcomitrella FtsZ1-2 in *E. coli*. BL21 Star™ (DE3) cells transformed with the constructs for eGFP-FtsZ1-2 were induced with 0.5 mM IPTG. Different elongated bacteria were observed (**A-E**). Filament formations are highlighted with arrows. Scale bars 5 µm.

**Figure S6.**
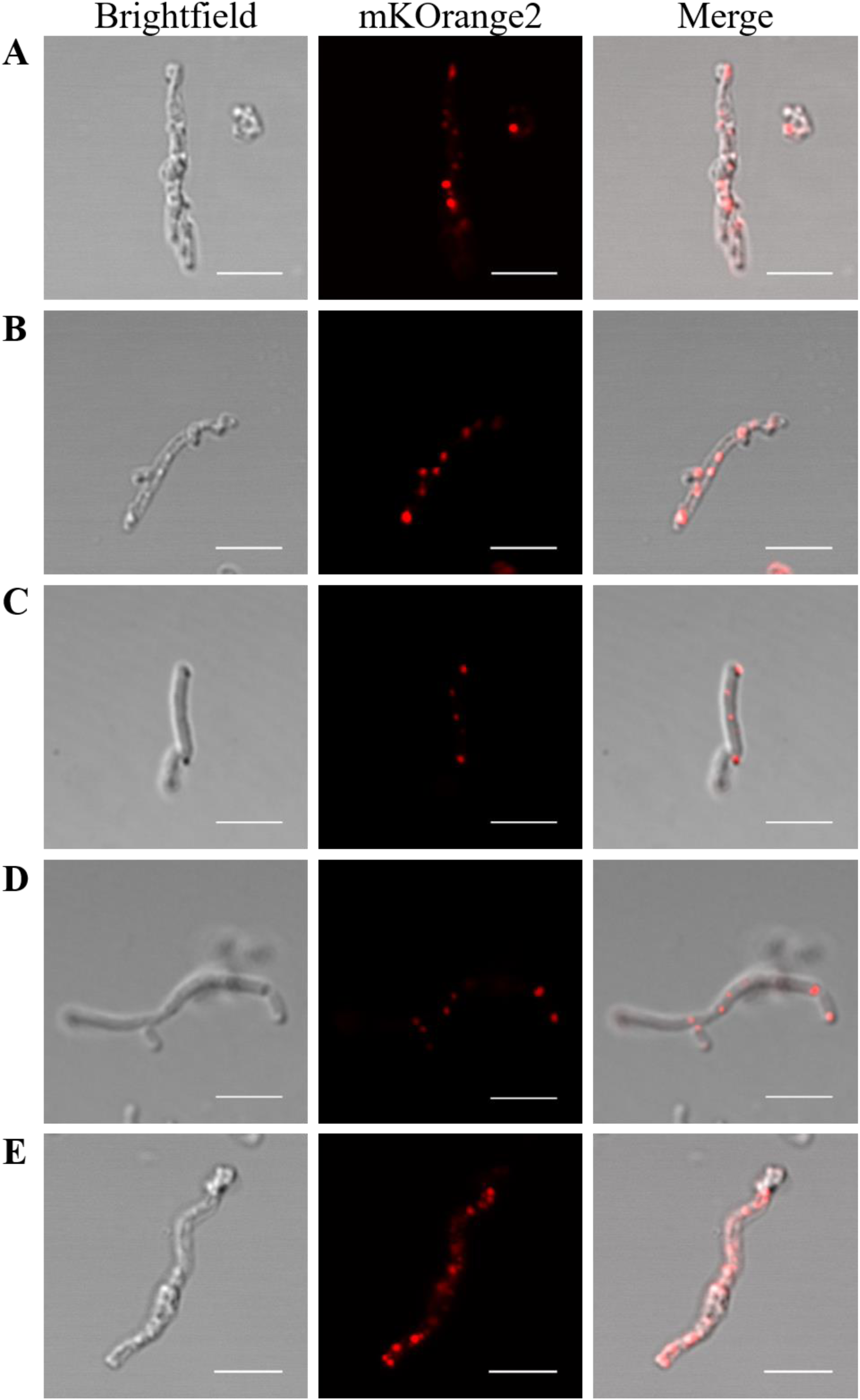
Confocal microscopy of mKO2-tagged Physcomitrella FtsZ2-1 in *E. coli*. BL21 Star™ (DE3) cells transformed with the constructs for mKO2-FtsZ2-1 were induced with 0.5 mM IPTG. Different elongated bacteria were observed (**A-E**). Scale bars 5 µm.

**Figure S7.**
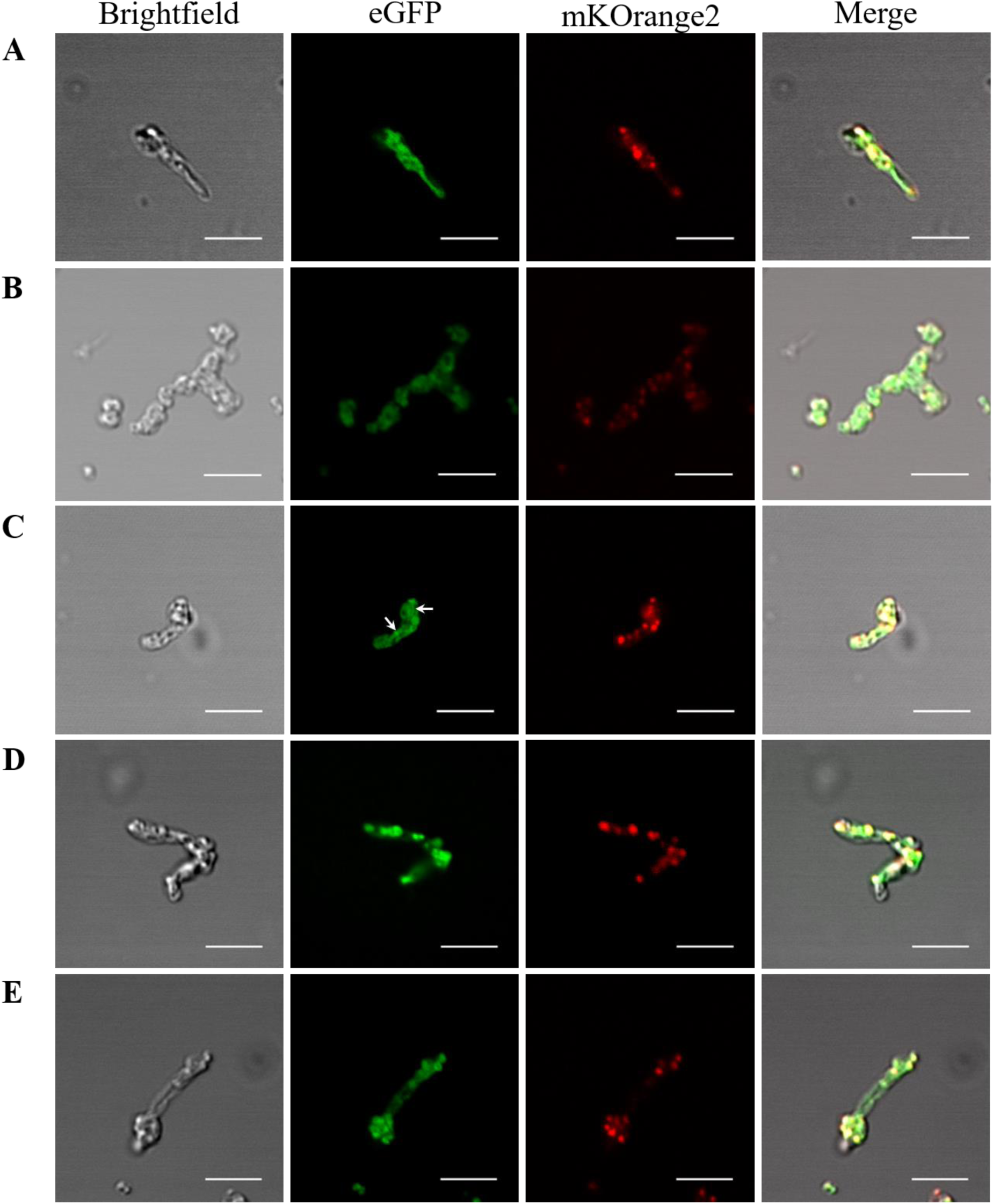
Confocal microscopy of eGFP-tagged Physcomitrella FtsZ1-2 and mKO2-tagged Physcomitrella FtsZ2-1 in *E. coli*. BL21 Star™ (DE3) cells transformed with the construct for eGFP-FtsZ1-2_mKO2-FtsZ2-1 were induced with 0.5 mM IPTG. Different elongated bacteria were observed (**A-E**). Filament formations are highlighted with arrows. Scale bars 5 µm.

**Figure S8.**
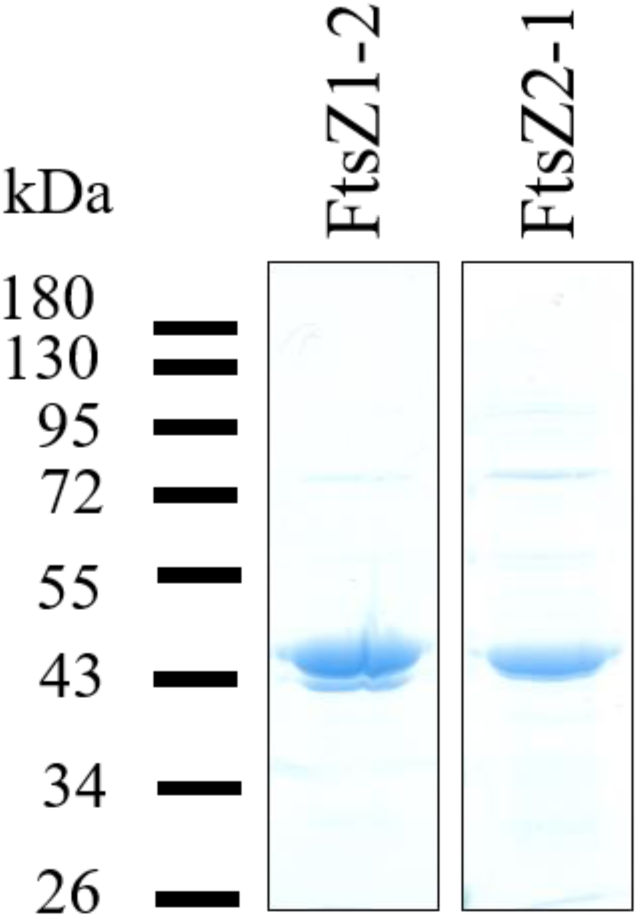
Coomassie-stained SDS-PAGE of purified FtsZ1-2 and FtsZ2-1. Physcomitrella FtsZ1-2 and FtsZ2-1 were overexpressed in *E. coli* as His₆-SUMO fusion proteins. The proteins were purified using Ni-affinity chromatography, followed by removal of the His₆-SUMO tag and further purification *via* size exclusion chromatography. The fractions obtained after size exclusion chromatography are displayed.

**Figure S9.**
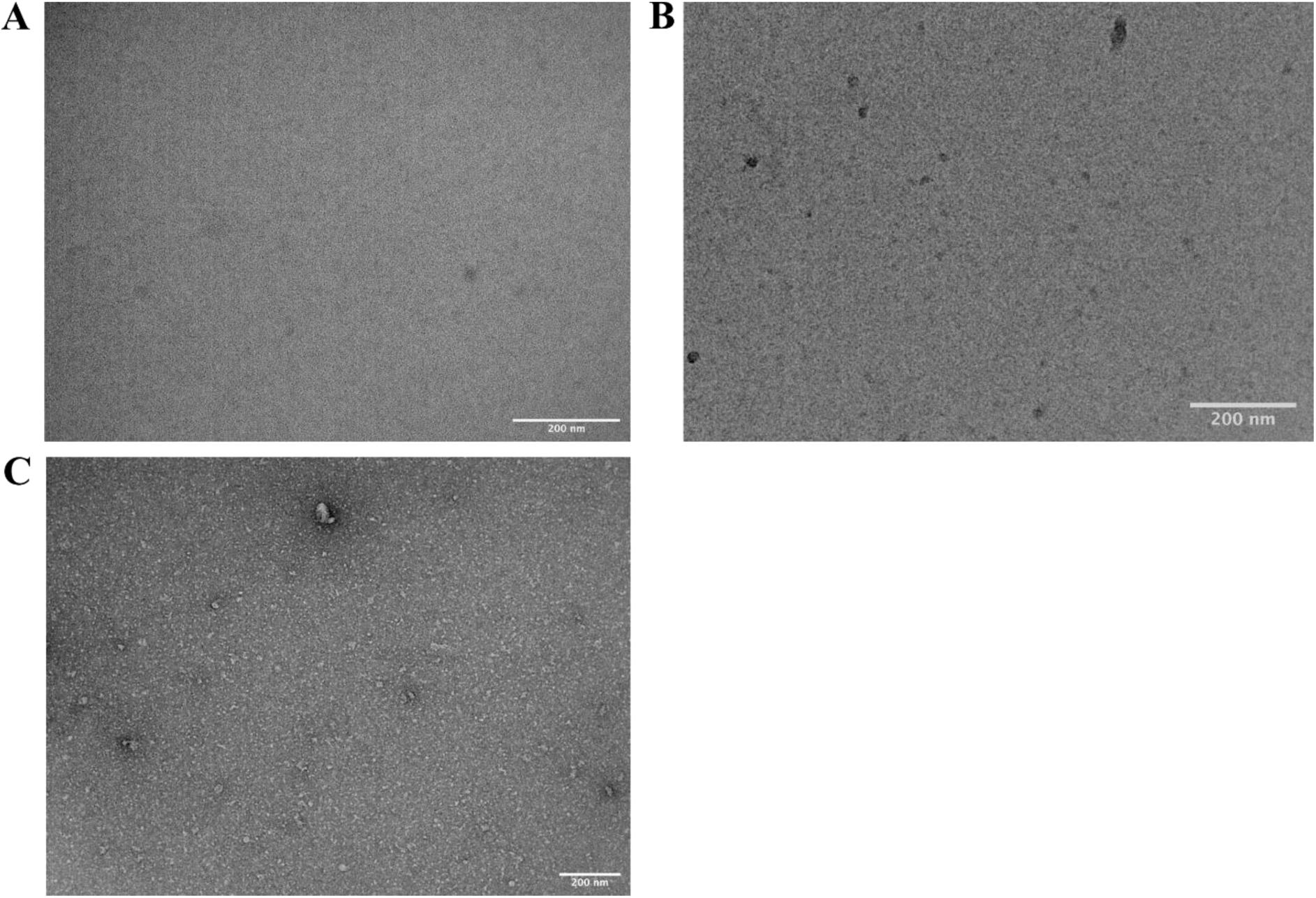
Negative stain transmission electron microscopy of Physcomitrella FtsZ1-2, Physcomitrella FtsZ2-1 together with FtsZ2-1 without GTP as negative control. 15 µM of FtsZ1-2 (A), 15 µM FtsZ2-1 (**B**) and 7.5 µM FtsZ1-2 mixed with 7.5 µM FtsZ2-1 (**C**) were incubated at room temperature for 5 min without GTP and then imaged by transmission electron microscopy. Representative images are shown. Scale bars as indicated. Experiments were repeated independently twice with similar results.

**Figure S10.**
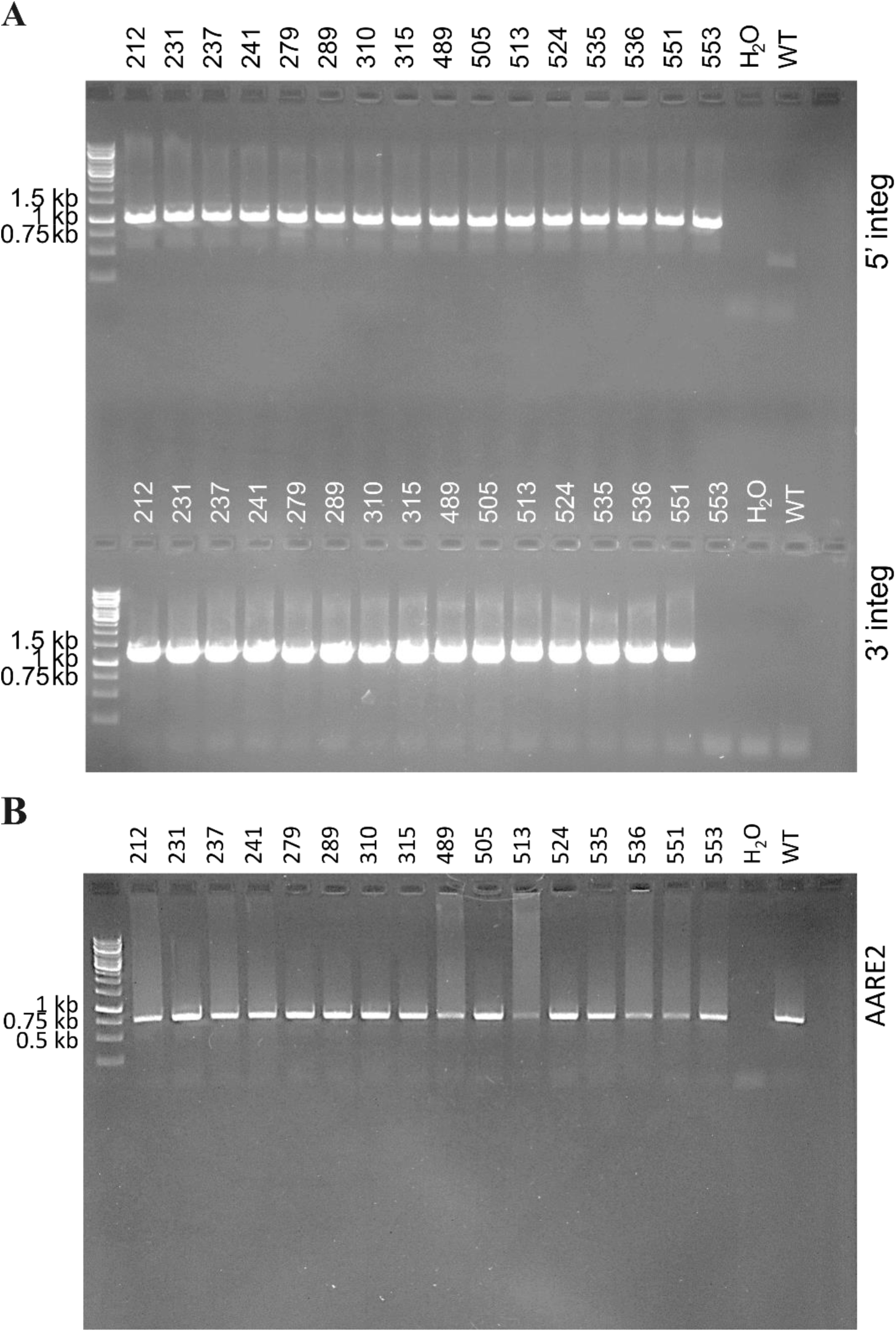
Uncropped gel image of the PCR result of the screening for transgenic FtsZ2-1-eGFP lines. (A) PCR testing correct integration at the selected 5’ and 3’ region. Expected amplicon sizes were 1064 bp (5’ integration) and 1302 bp (3’ integration). (B) Control PCR on AARE2 (Pp3c12_21080V3). Expected amplicon size was 768 bp. Primers for AARE2 were taken from [78].

## Notes

https://zenodo.org/records/10648779

### Competing Interest Statement

The authors have declared no competing interest.

### Summary of Updates

We optimized further our protein purification protocols. This resulted in more active Physcomitrella FtsZ proteins. We re-run all our in vitro assays with these highly active proteins and now provide consistent results. They show a synergistic interaction of the two Physcomitrella FtsZ isoforms between the different assays. Thus, we also improved the discussion. Reflecting the different work load for the revision, we changed the order of authors in four positions.

